# Mechanisms of Tone-in-Noise Encoding in the Inferior Colliculus

**DOI:** 10.1101/2025.03.07.642101

**Authors:** Johanna B. Fritzinger, Laurel H. Carney

## Abstract

Extracellular single-unit responses to tone-in-noise (TIN) stimuli were recorded in the inferior colliculus (IC) of awake female Dutch-belted rabbits. Stimuli consisted of wideband and narrowband tone-in-noise (TIN) with on-and off-characteristic frequency tones. Neural responses to wideband TIN showed a pattern of rates that increased when the tone matched CF and decreased (with respect to noise-alone responses) when the tone was above or below CF. This result differed from narrowband TIN IC responses that depended on envelope fluctuations in the stimulus, consistent with neural-fluctuation sensitivity. The WB-TIN responses could be fit with a difference-of-gaussians model that had narrow excitation and broad inhibition; responses to TIN could not be predicted by response-maps or spectrotemporal receptive fields. Responses to diotic and contralateral presentations of WB-TIN did not differ due to presentation ear. A single-CF computational model of the IC could not predict responses to wideband TIN. However, adding local off-CF inhibitory inputs to an on-CF IC model improved accuracy. These results suggest that broad inhibition could explain encoding of wideband TIN at suprathreshold signal-to-noise ratios, whereas neural fluctuation sensitivity is more important for narrowband sounds.

**Significance Statement:** Tone-in-noise (TIN) stimuli are used in physiology and psychophysics to explore sound perception. Physiological studies have primarily used tones at the characteristic frequency (CF) of a neuron. Here we present a systematic investigation of inferior colliculus (IC) neuron responses to on- and off-CF tones in wideband and narrowband noise, presented diotically or contralaterally, which revealed rate reductions not previously reported. Interestingly, responses to narrowband TIN can be explained by neural fluctuation sensitivity in the IC, but responses to wideband noise resulted in fundamentally different patterns of neural activity. Adding local off-CF inhibitory inputs to an established IC computational model accurately predicted responses to wideband TIN. This work highlights the role of broad inhibition in encoding complex sounds in the IC.

## 1. Introduction

In this study we compared mechanisms that influence tone-in-noise (TIN) encoding in the inferior colliculus (IC). TIN stimuli have been used extensively to gain insight on aspects of auditory perception, including energy and envelope cues, binaural perception, and critical bands (Fletcher, 1940; Kidd et al., 1989; Richards, 1992; van de Par and Kohlrausch, 1999). However, neural mechanisms underlying TIN representation in the IC are not well understood. Neural fluctuations (NFs) are hypothesized to provide a cue for tone detection in noise. NFs, or low-frequency changes in amplitude of auditory-nerve (AN) rate functions due to beating between stimulus components, are converted to rate changes by periodicity-tuned neurons in the IC (Carney, 2018, 2024). Addition of a tone to a noise reduces the depth of envelope fluctuations in the stimulus (Richards, 1992) and of NFs (Carney, 2018), providing a cue for tone detection. IC responses are consistent with NF predictions in awake rabbit for an on-characteristic frequency (CF) tone in a narrowband (1/3-octave) noise (Fan et al., 2021, 2022)(Fig. 1a). To further explore how NFs influence physiological responses to TIN, we recorded responses in the IC using a novel paradigm: a wideband noise with an added tone that was stepped past the CF (Fig. 1b). The predicted response to WB-TIN was a decrease in rate near- and on-CF, dependent on the periodicity tuning in the IC neuron (Fig. 1c,d). Interestingly, the qualitative trends in responses to wideband TIN (WB-TIN) differed from narrowband TIN (NB-TIN) responses, suggesting that mechanisms in addition to NFs are involved. This study investigated how NFs and inhibition interact to represent WB-TIN in the IC.

**Figure 1.**
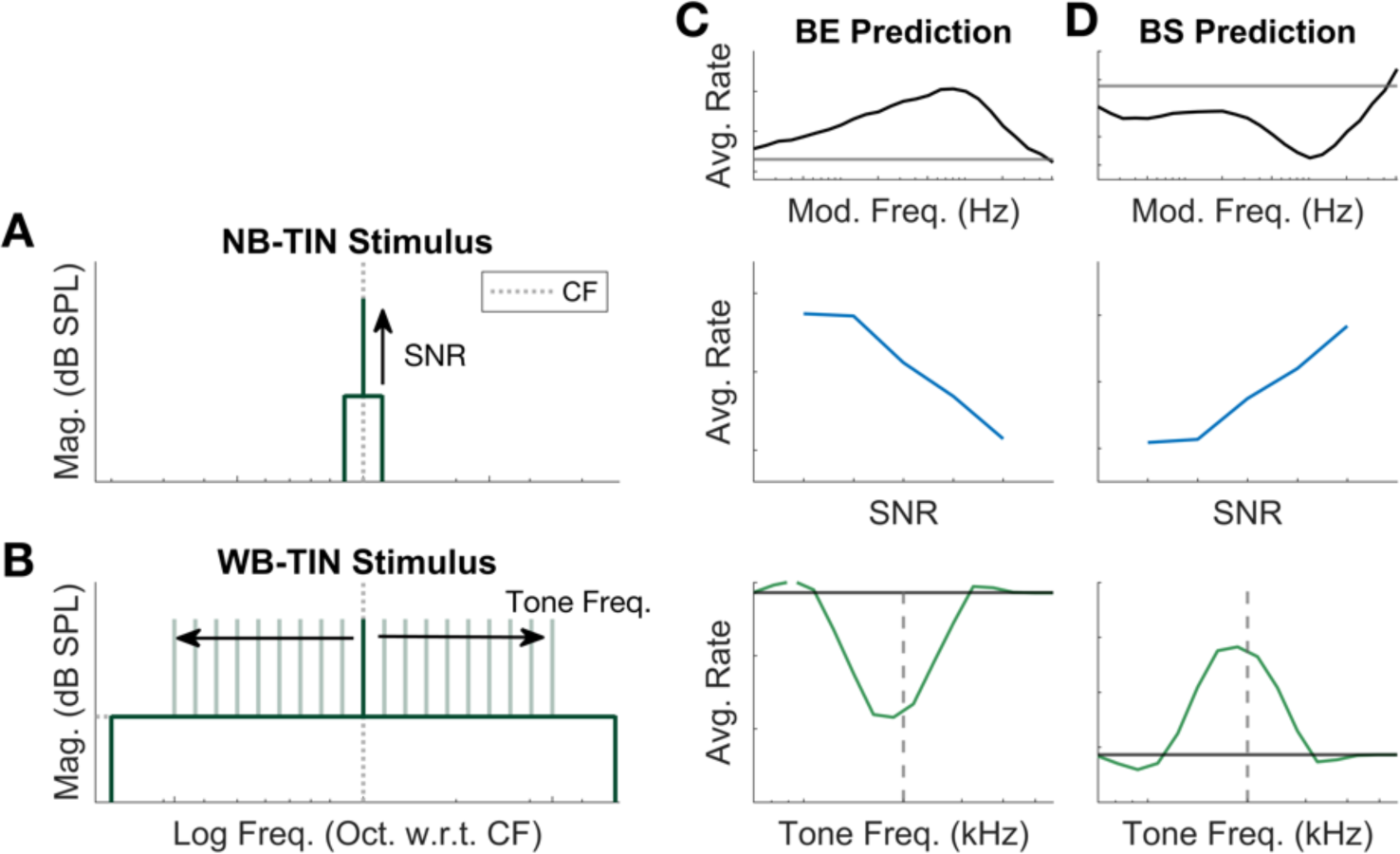
Illustration of stimuli and neural fluctuation hypothesis using a computational model of the IC. (A) NB-TIN stimulus with tone increasing in SNR at CF (B) WB-TIN stimulus with tone varying in frequency. (C) BE prediction based on neural fluctuation mechanism. MTF (top), response to increasing SNR NB-TIN (middle), and WB-TIN (bottom). (D) BS prediction based on neural fluctuation mechanisms, MTF, NB-TIN, and WB-TIN responses.

TIN stimuli have been widely used in psychophysical and physiological studies to investigate auditory filters in the context of the power-spectrum model, which postulates that tones are encoded by increased energy in the response of peripheral filters tuned to the tone frequency (Moore, 1975; Patterson, 1976). Physiological evidence for energy coding of on-CF tones in broadband noise include increasing rate as a function of signal-to-noise ratio (SNR) in IC neurons (Jiang et al., 1997; Rocchi and Ramachandran, 2018). However, the power-spectrum model is not consistent with psychophysical results, which show little effect on TIN detection performance when energy cues are made unreliable by random variation of sound level across intervals (Kidd et al., 1989; Richards, 1992). Psychophysical and modeling studies report that envelope-based cues are sufficient to describe TIN detection thresholds and outperform energy-based models (Dau et al., 1996; Kohlrausch et al., 1997; Richards, 1992; Davidson et al., 2009; Mao et al., 2013; Mao and Carney, 2015). These psychophysical and modeling TIN results suggest a role for the IC in representation of envelope cues because most IC neurons are sensitive to amplitude modulation (AM) (Langner and Schreiner, 1988; Krishna and Semple, 2000; Joris et al., 2004; Nelson and Carney, 2007; Kim et al., 2020).

Inhibition also plays an important role in shaping IC cell responses to sound (Casseday et al., 1994; Koch and Grothe, 1998; Caspary et al., 2008; Pollak et al., 2011), though the effect of inhibition on TIN responses has not been fully explored. Two-tone paradigms in the IC have revealed inhibitory response regions that may influence TIN encoding (Egorova et al., 2001; Portfors and Felix, 2005; Alkhatib et al., 2006). The frequency range over which neurons are excited broadens when inhibition is blocked or reduced, supporting a role of broad inhibition in shaping responses to tones (Yang et al., 1992; Palombi and Caspary, 1996; LeBeau et al., 2001; Xie et al., 2005). Sideband inhibition is observed in the response maps (RMs) of I-type IC neurons, which are hypothesized to encode tones in background noise levels more robustly than other RM types (Ramachandran et al., 1999, 2000).

This study investigated how NF sensitivity and off-CF inhibition contribute to TIN encoding by characterizing responses in the IC of awake rabbit to TIN stimuli with on- or off-CF tones. Responses of single neurons were compared for on-CF WB-TIN and NB-TIN. Next, responses to WB-TIN were characterized for a range of spectrum levels and SNRs, for contralateral and diotic stimuli. A difference-of-gaussians (DOG) model (Su and Delgutte, 2020) was used to explore excitatory and inhibitory components in the WB-TIN response. Lastly, computational IC models tested the hypothesis that interactions of NFs and off-CF inhibition influence TIN encoding.

## 2. Materials and Methods

### 2.1 Animals & Surgical Procedures

All procedures were approved by the University of Rochester Committee on Animal Resources in compliance with National Institutes of Health Guidelines. Four female Dutch-belted rabbits (*Oryctolagus cuniculus*) ranging from 6 months to 5 years of age were screened for normal hearing periodically using distortion product otoacoustic emissions (Whitehead et al., 1992), and experiments were discontinued in an animal if emission amplitudes decreased by 10 dB.

Surgeries included an initial headbar placement, initial craniotomy and microdrive placement, and microdrive replacements. For all surgeries, rabbits were anesthetized with either 66 mg/kg intramuscular ketamine and 2 mg/kg intramuscular xylazine or 35mg/kg intramuscular ketamine and 0.10-0.15mg/kg intramuscular dexmedetomidine. Animals were given the analgesic meloxicam (0.2mg/kg) subcutaneously for three days following surgery and were monitored for normal behavior.

A custom 3D-printed plastic headbar (ProtoLabs, Maple Plain, Minnesota) was affixed to the dorsal surface of the skull using stainless-steel screws and dental acrylic. After a one-month recovery period, the initial craniotomy and placement of the microdrive was performed. The microdrive was replaced every 1-6 months and the placement of the tetrodes was varied to sample different locations in the IC. Custom earmolds (Dreve Otoform Ak, Unna, Germany) were cast at the end of the microdrive surgeries while the animal was still under anesthesia.

For daily two-hour physiological recording sessions in a sound-attenuated booth (Acoustic Systems, Austin, Texas), awake rabbits were placed in a custom chair and the head was fixed. Custom earmolds were inserted and audio presentation hardware was calibrated at the beginning of the session using a probe-tube microphone (Etymotic ER10B+ or ER7C, Etymotic Research, Inc., Elk Grove Village, Illinois).

### 2.3 Microdrive and Electrodes

Extracellular recordings were made using a chronically implanted microdrive. The microdrive was 3D printed plastic, modeled after the Neuralynx Five-drive microdrives and supporting EIB-16 Five-Drive board connectors (Neuralynx, Inc., Bozeman, MT). The microdrive positioned four tetrodes, which were made by twisting four 18-μm platinum-iridium wires insulated with epoxy (California Fine Wire Co., Grover Beach, CA). The tetrodes were inserted into polyimide tubes (Polymicro Technologies Inc., Phoenix, AZ) which were passed through four stainless-steel 28-gauge guide tubes (Eagle Stainless, Franklin, MA). Tetrodes were plated with platinum-black plating solution (Neuralynx, Inc., Bozeman, MT) to lower impedances to a range from 0.1-1.5 MΩ.

### 2.4 Spike Sorting & Clustering

Signals were recorded using the Intan RHD recording system, including a 16-channel headstage with initial filtering stages (16-Channel Headstage with RHD2132 chip, Intan Technologies, LLC., Los Angeles, CA). The sampling frequency was set at 30 kHz. These initial filters included a 1^st^-order, analog, high-pass filter (Fc = 150 Hz), a 3^rd^-order, analog, Butterworth low-pass filter (Fc = 7.5 kHz), and a digital high-pass filter (Fc = 300 Hz) and saved by the Intan RHD recording system for post-hoc filtering and analysis (Intan Technologies, LLC., Los Angeles, CA).

To isolate neurons for analysis, signals were filtered, thresholded, and then clustered into separate neurons, as described below. First, the signals were filtered with a 4^th^-order Butterworth bandpass filter (300-3000 Hz). The majority of potential spikes were detected using a threshold of 4 times an estimate of the standard deviation (STD), with the standard deviation estimated by 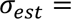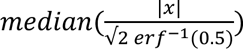 (Quiroga et al., 2004). The threshold was adjusted higher for a few sessions to improve clustering of neurons that appeared to be separable but were not clustered well using the 4 x STD threshold criterion. For a few sessions, artifacts due to movement were present during the first dataset, and a different dataset was used to calculate the STD of the signal. If two above-threshold events occurred within 0.5 ms, only the event with the larger amplitude was determined to be a potential spike. All potential spike waveforms and times were then saved for clustering. Most neurons were clustered by comparing repolarization slopes; a few neurons were better isolated using principal components analysis. <u>(Schwarz et al., 2012</u>)

Three criteria were required to classify a recording as a well-isolated neuron. First, less than 2% of the inter-spike intervals could be less than 1-ms. Second, spike waveforms for each wire, when aligned by their peaks, were required to have similar shapes. Third, although many recordings could potentially have been clustered into several neurons, clustering was deemed valid only if the quantitative cluster-separation metric (Schwarz et al., 2012) was < 0.1. Cells were classified as unique neurons when the tetrodes were moved and the response properties of the neuron changed.

To verify that tetrode locations were in the central nucleus of the IC, responses to a set of characterizing stimuli were monitored for properties that matched typical IC recordings, e.g., responses to tones and diotic or dichotic noises. CFs were also monitored to check that as tetrode depth increased, CF increased (Schreiner and Langner, 1997). After an animal showed a 10-dB reduction in emission amplitudes or had elevated neural thresholds, the animal was euthanized and histology was completed to confirm tetrode placement.

### 2.5 Basic Stimuli & Analysis

Stimuli were created in MATLAB (Mathworks, Natick, MA) and presented through an audio interface (16A, Mark of the Unicorn, Cambridge, Massachusetts), a digital-to-analog converter (DAC3 HGC, Benchmark Media Systems, Inc., Syracuse, New York), and earphones (Beyerdynamic DT-48, Beyerdynamic GmbH and Co., Heilbronn, Germany or Etymotic ER2, Etymotic Research, Inc., Elk Grove Village, Illinois). The earphones were coupled with the custom earmolds. The sampling frequency for sound presentation was 48,000 Hz.

At the beginning of each session, a set of stimuli was presented to characterize the neuron. These stimuli included tones to construct an RM, sinusoidally modulated noise for a modulation transfer function (MTF), binaural noise to characterize basic binaural response properties, and wideband noise to construct a spectrotemporal receptive field (STRF).

The RM was created by presenting pure-tone stimuli at 10, 30, 50, and 70 dB SPL, with frequencies spanning 250-16000 Hz, with five steps per octave. The stimuli were diotic, 200 ms in duration, with 10-ms cos^2^ on/off ramps, repeated 3 times in a random sequence. The CF was estimated by finding the tone frequency that elicited a response at the lowest sound level. If the CF based on the response to pure tones was unclear, the peak of the noise STRF was used to estimate CF. Neurons were categorized as V-type, I-type, O-type, onset, onset/offset, inhibitory, and unusual. Only V-type, I-type, and onset neurons are included in analysis due to small sample size of other categories. Onset neurons were defined as neurons with an onset response but no sustained response to the RM stimulus. I-type neurons had a sustained response and the bandwidth at 70 dB SPL did not exceed 2 times the bandwidth at threshold. Neurons were categorized as V-type if they had a sustained response and the bandwidth at 70 dB SPL was greater than 2 times threshold bandwidth. For some analyses, neurons were separated into three groups based on CF: low (CF < 2 kHz), medium (2 kHz ≤ CF < 4 kHz), and high (CF ≥ 4 kHz).

The MTF of the neuron was determined by presenting 100% sinusoidally amplitude-modulated broadband (100-10000 Hz) diotic, flat-spectrum, gaussian noise. The amplitude-modulation frequencies tested were 2-600 Hz in increments of three steps per octave, presented at 33 dB SPL spectrum level, with a duration of 1 s including 50-ms cos^2^ ramps, and repeated five times, in a random sequence. Each neuron was categorized into one of four MTF classification types: band-enhanced (BE), band-suppressed (BS), hybrid, and flat (Kim et al., 2020). MTFs were classified as BE if at least two average-rate responses to modulation frequencies were significantly larger than the unmodulated rate (i.e., average rate in response to an unmodulated noise), over a range of modulation frequencies that was not interrupted by a rate significantly lower than the unmodulated rate (unpaired t-test, p < 0.05)(Fig. 1c). Conversely, MTFs were classified as BS if at least two average-rate responses to modulation frequencies were significantly lower than the unmodulated rate, over a range of modulation frequencies that was not interrupted by a rate significantly above the unmodulated rate (unpaired t-test, p < 0.05)(Fig. 1d). A cell was classified as hybrid if conditions for both BE and BS categories were met. A cell was classified as flat if neither BE nor BS condition was met. The best modulation frequency (BMF) for BE and hybrid neurons was the modulation frequency that resulted in the highest spline-interpolated, average-rate response. Conversely, the worst modulation frequency (WMF) for BS and hybrid neurons was the modulation frequency that resulted in the lowest spline-interpolated average-rate response. Hybrid neurons were divided into two categories: neurons were classified as H_BE_ if the rate of the neuron in response to modulation frequencies near 100 Hz was greater than the unmodulated response and H_BS_ if responses near 100 Hz were suppressed. The responses near 100-Hz modulation were used in these categorizations because the hybrid neuron response properties were more similar to the BE or BS MTF-classification type that matched their response to 100-Hz modulation.

Binaural response properties were determined by presenting a wideband (100-15,000 Hz), flat-spectrum, gaussian noise at 0, 10, 20, or 30 dB SPL spectrum level to the ipsilateral, contralateral, or both ears. The stimuli were 1-s duration with 10-ms cos^2^ ramps and were presented three times in a random sequence. The average-rate responses were used to identify the contribution of each ear to the binaural response.

The STRF was estimated based on responses to flat-spectrum gaussian noise (100-16,000 Hz) with a 2-s duration, presented at an overall level of 68 dB SPL. Ten repetitions each of 25 different noise tokens were presented in a random sequence, either diotically or to the contralateral ear. The STRFs were created using a 2^nd^-order Wiener-kernel analysis based on Lewis et al. (2002). Briefly, the 2^nd^-order reverse correlation was computed between the gaussian-noise stimulus and the response of the neuron. Then, singular-value decomposition was used to decompose the kernel into excitatory and inhibitory components (Lewis et al., 2002).

### 2.6 Tone-in-Noise Stimuli & Analyses

The WB-TIN stimulus consisted of a broadband, flat-spectrum, gaussian noise (3- or 4-octave bandwidth) geometrically centered on CF with an added pure tone (Fig. 1b). The tone was varied in frequency over 3 octaves centered on CF, in steps of 6 per octave. SNRs tested were −inf (noise alone), 20, 30, and 40 dB tone level with respect to noise spectrum level in dB SPL. The SNRs were selected to straddle rabbit WB-TIN behavioral detection thresholds (Zheng et al., 2002). Stimuli were presented in random order, with 300-ms duration and 10-ms cos^2^ on/off ramps, and repeated 30 times, with each repetition being a different gaussian-noise token. Diotic and contralateral presentations of the WB-TIN stimulus had noise spectrum levels (N_0_) of 3, 23, or 43 dB SPL.

The NB-TIN stimulus was similar to the WB-TIN stimulus, with a few key differences (Fig. 1a). The tone frequency was shifted one octave above and one octave below CF, in 6 steps per octave. To create the NB-TIN stimuli, a single frozen broadband, flat-spectrum, gaussian noise was filtered to a 1/3-octave bandwidth centered on each tone frequency, using a 5000-th order finite-impulse-response filter. Spectrum level, SNR, duration, and ramp duration of the NB-TIN stimulus were matched to the WB-TIN stimulus. Twenty repetitions of each NB-TIN stimulus were presented; stimuli were diotic or contralateral. During recording sessions, the CF of the neuron was not precisely known, so shifting the NB-TIN spectrum in frequency ensured that the neural response to an on-CF tone was recorded.

The rate profiles for WB-TIN and NB-TIN were calculated by averaging the firing rate over the duration of the stimulus, excluding a 50-ms onset, and plotted as a function of tone frequency. On-CF responses for NB- and WB-TIN were determined by taking the average rate response of the neuron for the stimulus with the tone nearest to CF. The proportion of predictable variance was quantified by V_p_ = 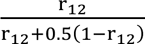, where *V_p_* is the proportion of predictable variance, and 𝑟_12_ is the first-half ̶ last-half correlation between responses to repetitions of the stimulus (Ahumada and Lovell, 1971).

WB-TIN rate profiles were compared to linear predictions based on RMs and noise STRFs. Variance explained was calculated between the RM level closest to the overall level of the WB-TIN and the WB-TIN response. To compare WB-TIN responses to the noise STRF of the neuron, the noise STRF was used to predict a neuron’s response to the WB-TIN stimulus. First the spectrogram of the sound stimulus was calculated. Then the spectrogram was convolved frequency-by-frequency with the noise STRF. Next, the result was summed over frequency to create a predicted peri-stimulus time histogram (PSTH), which was then averaged over time to compute a predicted average rate. The variance explained by the STRF-model average rate was calculated to determine similarity between STRF and WB-TIN responses.

Neural responses to WB-TIN were fit to a DoG function for further analysis of excitation and inhibition (Su and Delgutte, 2020):

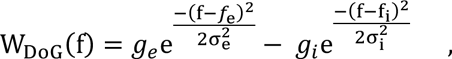

where *f* is the frequency range of the tone in the WB-TIN stimulus. The mean of the excitatory and inhibitory gaussian functions, 𝑓_𝑒_ and 𝑓_𝑖_, respectively, were near the CF of the neuron. All frequencies and bandwidths were fit in logarithmic units and transformed into linear units for plotting. The standard deviations, σ_𝑒_and σ_𝑖_, represent the excitatory and inhibitory bandwidths. The strength of the excitatory and inhibitory gaussians are 𝑔_𝑒_ and 𝑔_𝑖_, respectively. The DoG function was fit to the neural data with the noise-alone response subtracted by minimizing the squared error using the MATLAB function fmincon.

### 2.7 Statistical Analyses

Two linear mixed-effect models were fit to detect statistically significant changes in the WB-TIN response due to stimulus parameters and neuron characteristics. MATLAB functions *LME* and *compare* were used for model selection and the statistical software package JASP (JASP Team, 2024, Version 0.18.3, computer software) was used for the remaining analysis. The WB-TIN rates for each neuron were normalized using a noise-referenced z-score approach: 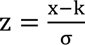, where *x* was the average rate response, *k* is the noise-alone rate, and *σ* is the standard deviation of *x*. Both models included neuron identity as a random effect, specifically a random intercept, to account for the differences in rate for each neuron.

The first model included the following stimulus parameters as fixed effects: tone frequency, in −1 to 1 octaves above CF, in increments of 0.25 octaves, spectrum level, SNR, and presented ear. All fixed effects were coded as categorical variables. The model chosen for this analysis included two-, three-, and four-way interactions between fixed effects.

The second model included the following stimulus parameters and neural characteristics as fixed effects: tone frequency, spectrum level, CF group (low, medium, high), MTF type, and RM type as categorical variables. The data for this second model was curated to include 40-dB SNR data and only V- and I-RM types due to small sample sizes of other RM types. The binaural/contra variable did not improve model fit and was not included as a parameter. The model chosen for this analysis included two-, three-, and four-way interactions of the fixed effects. Both model structures were selected using forward stepwise regression based on the Akaike information criterion.

The model fits were assessed by visual examination of residual and quantile-quantile plots in MATLAB. The models were analyzed using type III ANOVAs in JASP using the Satterthwaite approximation. Further analysis of the results was done using contrast testing. Other statistical methods used throughout this study include t-tests, ANOVAs, and comparisons of correlation coefficients and variance explained.

### 2.8 Modeling

All models used the WB-TIN stimulus with a sampling rate of 100 kHz. To compare the neuron responses to WB-TIN to predictions of the energy model, an estimate of energy was computed by passing the stimulus through a 4^th^-order gammatone filter and taking the RMS of the filter response. Each gammatone filter was centered on the CF of the neuron of interest. The gammatone tau parameter was calculated by finding the cat Q_10_ value for a specific CF, (Q_10_ = 10^0.4708∗log10(CF/1000)^ + 0.4664) (Carney and Yin, 1988), converting to ERB_CF_ = CF/Q_10_, then calculating tau, 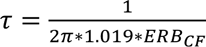 (Patterson et al., 1988).

For all remaining models, a phenomenological AN model stage (Zilany et al., 2014) was followed by a same-frequency inhibition-excitation (SFIE) IC model (Nelson and Carney, 2004; Carney and McDonough, 2019). The SFIE model is a phenomenological model that replicates amplitude-modulation sensitivity seen in physiological recordings of IC neurons, including BE (Fig. 2c) and BS (Fig. 2d) MTFs (Joris et al., 2004; Nelson and Carney, 2004; Kim et al., 2020). The IC model was expanded to include three SFIE paths with different CFs: one centered on the tone frequency, and two off-CF pathways symmetrically in octaves above and below the on-CF pathway. Six different configurations with off-CF inhibition were tested: ascending off-CF inhibition from cochlear nucleus (CN) to a BE or BS cell, lateral off-CF inhibition from a BS or BE off-CF cell to a target on-CF BE cell, and lateral off-CF inhibition from a BE or BS cell to a target on-CF BS cell (Fig. 2c).

**Figure 2.**
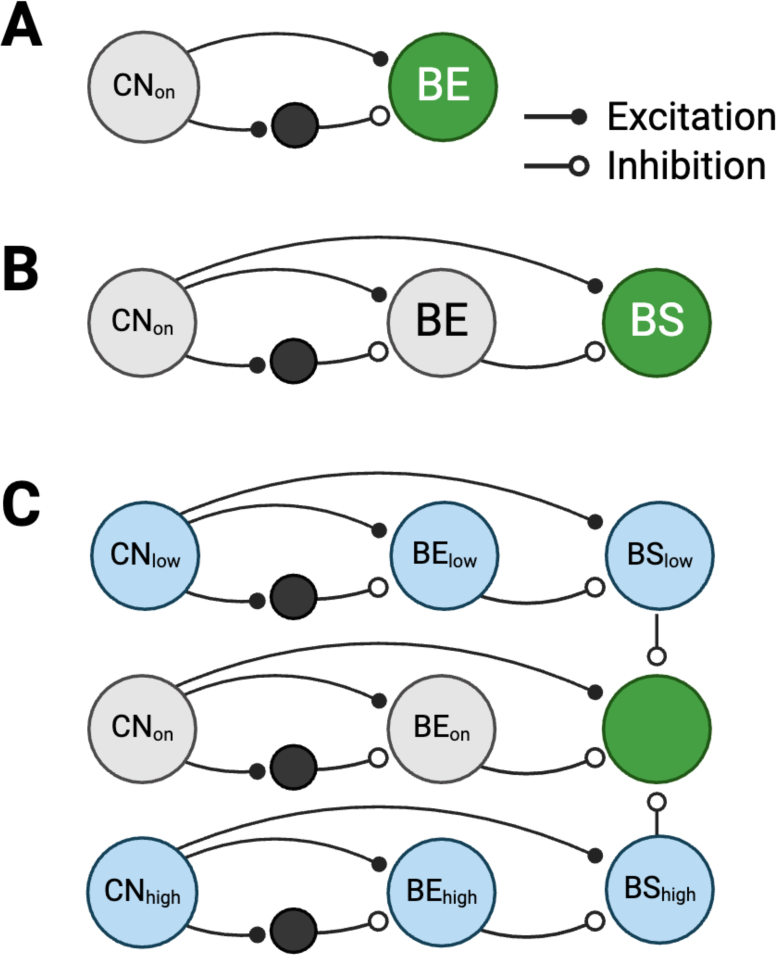
Schematics of SFIE IC model and one configuration that added off-CF inhibition. (A) SFIE model configuration with an on-CF CN stage (CN_on_) and IC BE output cell (green). (B) SFIE model with an output cell that is an IC BS cell (green). (C) Model configuration with the output model cell as a BS on-CF SFIE BS cell inhibited by off-CF BS cells.

All model configurations had on-CF excitation, on-CF inhibition, and two off-CF inhibitions, based on this structure:

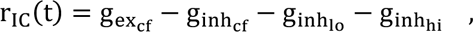

where g_exc𝐹_ was the excitatory on-CF input, g_inhc𝐹_ was the inhibitory on-CF input, g_inhlo_ was the inhibitory input from below CF, and g_inhhi_ was the inhibitory input from above CF. The combined response, 𝑟_𝐼𝐶_(𝑡), which represented the instantaneous firing rate of the IC model neuron, was half-wave rectified. The four model inputs were described as

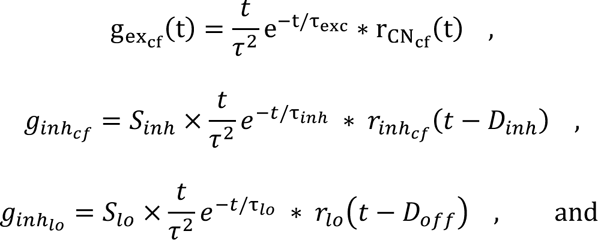

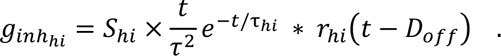

Each equation represents an input convolved with an inhibitory or excitatory alpha function, where the alpha function was normalized to an area of 1. The inhibitions were delayed, by *D*, and multiplied by a strength term, *S.* The excitation strength was 1. The input 𝑟_𝐶𝑁𝑐𝑓_ is the instantaneous rate of the cochlear nucleus on-CF model neuron, the inputs 𝑟_𝑖𝑛ℎ𝑐𝑓_, 𝑟_𝑙𝑜_, and 𝑟_ℎ𝑖_, and 𝑆_𝑖𝑛ℎ_ and 𝐷_𝑖𝑛ℎ_, the strength and delay of the on-CF inhibition, respectively, varied across the six model configurations (Table 1). The time constants, 𝜏, were parameters for the alpha functions, where 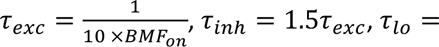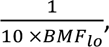 and 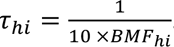.

**Table 1.** Model inputs and fixed parameters for all six lateral inhibition model configurations.

| | $r_{inh_{CF}}$ | $r_{lo}$ | $r_{hi}$ | $S_{inh}$ | $D_{inh}$ |
| --- | --- | --- | --- | --- | --- |
| BE inhibited by off-CF CN | $r_{CN_{cf}}$ | $r_{CN_{lo}}$ | $r_{CN_{hi}}$ | 0.9 | $2\tau_{exc}$ |
| BE inhibited by off-CF BE | | $r_{BE_{lo}}$ | $r_{BE_{hi}}$ | | |
| BE inhibited by off-CF BS | | $r_{BS_{lo}}$ | $r_{BS_{hi}}$ | | |
| BS inhibited by off-CF CN | $r_{BE_{cf}}$ | $r_{CN_{lo}}$ | $r_{CN_{hi}}$ | 4 | $2\tau_{exc}$ |
| BS inhibited by off-CF BS | | $r_{BS_{lo}}$ | $r_{BS_{hi}}$ | | |
| BS inhibited by off-CF BE | | $r_{BE_{lo}}$ | $r_{BE_{hi}}$ | | |

The parameters that were fit to each model configuration were the off-CF values, 𝑆_𝑙𝑜_, 𝑆_ℎ𝑖_, and BMF_on_, BMF_lo_, and BMF_hi_. The delay parameter, 𝐷_𝑜𝑓𝑓_, was set to 1 ms after an initial grid search revealed that the delay parameter did not impact average-rate results. The stimuli used to fit the model to example neural data included the MTF, an on-CF NB-TIN at 23 dB SPL and −inf, 20, 40 dB SNR, and a WB-TIN stimulus at N_0_ = 23 dB and 40 dB SNR. For the fitting procedure, first the AN model was run with three different CFs: CF_lo_, CF_on_, and CF_hi_, where the low and high CF value was varied in increments of 1/6 octave +/− CF up to 1.5 octaves +/− CF. Then, the MATLAB function fmincon with the sequential quadratic programming (SQP) algorithm was used to minimize an objective function to fit the broad inhibition parameters for each CF range. The fit parameters were initialized with random numbers within a bounded range. The range for strength parameters was 0 to 1, and the range for BMFs was 10 to 300 Hz. The objective function was 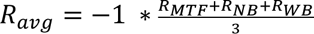, where R_avg_ is the mean of the correlation between the data and the predicted model for each dataset. After a model was fit to each CF range (12 total), the CF range and parameter set with the highest average correlation coefficient was used as the best fit to the data. CF range was not included in the gradient descent method due to the computational expense of running the AN model for each CF.

#### Code Accessibility

Custom MATLAB code for clustering data (Schwarz et al., 2012) available at https://www.urmc.rochester.edu/labs/carney/publications-code/spike-sorting-code.aspx. Custom MATLAB code for data analysis and modeling along with data files are available at https://osf.io/p4r82/. Code is also available on GitHub at https://github.com/jfritzinger/FritzingerCarney2025-WBTIN.

## 3. Results

Single-neuron activity was recorded from 229 neurons in the central nucleus of the IC in four awake Dutch-belted rabbits. All neurons had a response to binaural broadband noise greater than 5 sp/s. Responses of 127 neurons to both diotic and contralateral WB-TIN were recorded. Responses of an additional 80 neurons to diotic WB-TIN, and 21 neurons to contralateral WB-TIN were recorded. Responses to NB-TIN were recorded in 106 of the neurons, and noise STRFs were recorded in 170 neurons. Distributions of neuron characteristics matched similar datasets in awake rabbit (Kim et al., 2020; Fan et al., 2021). The CFs ranged from 328 Hz to 10.6 kHz, with a median of 2.8 kHz (Fig. S1b). The MTF distribution was 58 BE (25.3%), 104 BS (45.4%), 43 hybrid (18.8%), and 24 flat (10.5%) (Fig. S1a). The BMF of the BE neurons ranged from 12 Hz to 412 Hz, with a median of 70 Hz (Fig. S1a). The worst modulation frequency (WMF) of BS neurons ranged from 2 Hz to 512 Hz with a median of 102 Hz (Fig. S1c). Hybrid neurons have both a BMF and a WMF; Hybrid BMFs ranged from 5 Hz to 512 Hz with a median of 64 Hz, and WMFs ranged from 2 Hz to 478 Hz with a median of 125 Hz (Fig. S1c). For the purpose of analysis and modeling, the predictable variance, V_p_, of the response to WB-TIN was calculated for all conditions to identify reliable responses; conditions with a V_p_ < 0.4 were excluded from analysis (Fig. S1d).

### Single-neuron response to frequency-shifted WB-TIN

IC neurons were initially characterized by a RM, MTF, and STRF (Fig. 3a, b). The noise STRF for the example in Fig. 3 featured early and later broad excitation centered on CF and both a lagging off- and on-CF inhibition (Fig. 3c). The rate in response to on-CF NB-TIN decreased with increasing SNR for two out of the three spectrum-level conditions (p<0.001 for N_0_ = 23, 43 dB SPL, p=0.400 for N_0_ = 3 dB SPL, t-test, Fig. 3d). This decrease in rate is consistent with a report of BE responses to on-CF tones in narrowband gaussian noise (Fan et al., 2021). In contrast, the response of this neuron to on-CF WB-TIN increased as SNR increased (Fig. 3h). Looking over a range of tone frequencies, in comparison to the noise-alone response (Fig. 3e-g, solid grey line), rates increased when the tone was near CF, and decreased for tones below or above CF (Fig. 3e-g). The excitation and flanking inhibitions were present at all three noise levels tested and excitation increased in magnitude as level increased. The difference between on-CF tones in wideband and narrowband noise was not expected in the neural fluctuation hypothesis, and off-CF rate decreases for WB-TIN were also not predicted.

**Figure 3.**
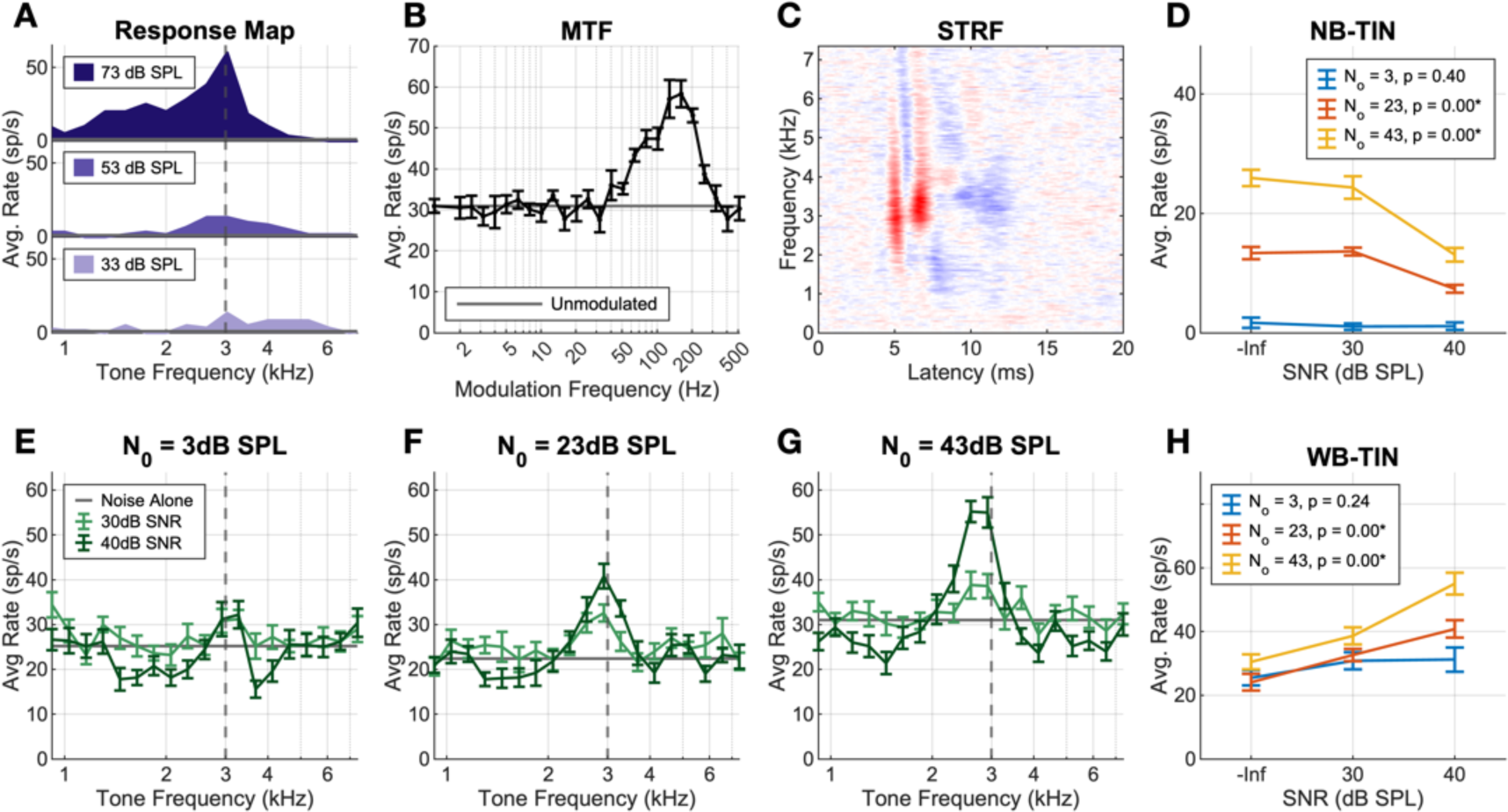
Single neuron responses to tones and noise used to characterize the neurons. (A) RM, dashed grey line represents estimated CF. (B) MTF, BE. (C) STRF estimated from responses to wideband noise, excitation in red and inhibition in blue. (D) Response to a NB-TIN centered on CF, for three levels (N_0_ = 3, 23, 43 dB SPL, overall level = 31, 51, 71 dB SPL) and 3 SNRs (−inf, 30, 40 dB). (E-G) Response to WB-TIN (N_0_ = 3, 23, 43 dB SPL, overall level = 43, 63, 83 dB SPL). Tones were presented at 30 and 40 dB SNR w.r.t. N_0_. The grey line is the noise-alone response. (H) Response to WB-TIN with tone at CF, rates estimated from data in (E-G). All error bars represent +/− 1 SEM.

### Responses to on-CF tones in WB and NB noise

To quantify differences in neural responses due to noise bandwidth when a tone was at CF, we compared responses to on-CF WB- and NB-TIN. Due to the previous finding that responses to NB-TIN are dependent on MTF type, neurons were divided into four categories, BE, BS, H_BE_, and H_BS_. An on-CF WB-TIN elicited an increase in rate as SNR increased for most neurons, regardless of MTF type, spectrum level, or contralateral or diotic presentation (Fig. 4). Rates for diotic presentations of the on-CF WB-TIN increased with SNR (n=57/84 N_0_=3, n=67/86 N_0_=23, 68/86 N_0_=43 dB SPL, Fig. 4a-c). In contralateral presentations, rate increased as SNR increased for the majority of neurons in all conditions (n=25/40 N0=3, n=30/42 N0=23, n=35/42 N0=43 dB SPL, S2). The contralateral 3-dB-SPL N_0_ condition was the only condition for which BE and BS cells had significantly different rate changes in the WB-TIN response (unpaired t-test, t(19)=-2.9263, p=0.0087, Fig. S2a). Overall, five out of six conditions did not show any effect of MTF type on responses to on-CF WB-TIN.

**Figure 4.**
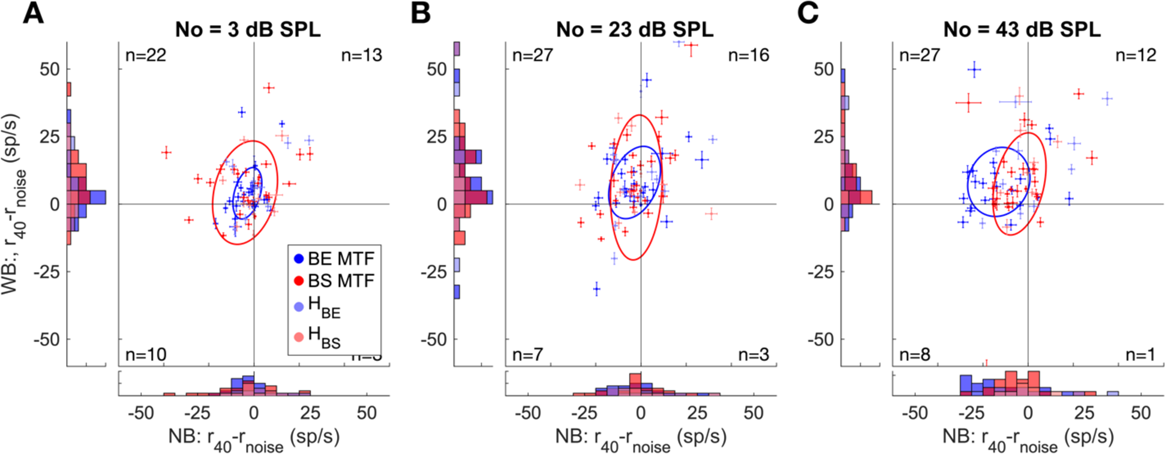
Differences between average rates in response to diotic TIN (40 dB SNR) and noise-alone for on-CF tones in wideband vs. narrowband noise. Responses at (A) N_0_ = 3 dB SPL, WB-TIN overall level range of 34-48 dB SPL, NB-TIN level range of 28-41 dB SPL, (B) 23 dB SPL, WB-TIN overall level range of 54-69 dB SPL, NB-TIN level range of 48-63 dB SPL, and (C) 43 dB SPL, WB-TIN overall level range of 74-83 dB SPL, NB-TIN level range of 68-83 dB SPL. BE MTFs are blue, BS MTFs are red, H_BE_ are light blue, H_BS_ are pink. Error bars represent +/− 1 SEM. Ellipses represent the standard deviation for BE (blue) and BS (red) neurons. Number of neurons in each quadrant is reported in the corner of each quadrant.

For the NB-TIN condition, responses of BE and BS neurons differed for the contralateral 23-dB-SPL N_0_ condition (unpaired t-test, t(25)=-3.1598, p=0.0041, Fig. S2b), and diotic 43-dB-SPL N_0_ condition (unpaired t-test, t(46)=-2.4818, p=0.0168, Fig. 4c). In both conditions, rates of BE cells decreased when the tone was added. BS cells in the contralateral 23-dB-SPL N_0_ condition had increased rates when the tone was added, and BS cells in the diotic 43-dB-SPL N_0_ condition had rates that either increased or decreased when the tone was added. The BE and BS responses were similar in the other four conditions, with distributions centered around 0, indicating that rate increases and decreases occurred in both BE and BS neurons. These NB-TIN on-CF results were only partially consistent with Fan et al. (2021), which reported MTF-type-dependent responses for many levels and SNRs. Potential explanations for the differences between studies are discussed below.

### Characterizing responses to frequency-shifted WB-TIN

The previous analysis found that most neurons responded to WB-TIN with an increase in rate at CF, and our next step was to determine how neurons responded to the rest of the WB-TIN stimulus, including for different sound levels and SNRs. A linear mixed-effects model was implemented to assess how stimulus parameters influence responses to WB-TIN. The mixed model revealed a significant three-way interaction between SNR, N_0_, and tone frequency (F_(32, 20580)_ =3.74, p=4.87e-12). SNR increases led to stronger rate increases near CF (+/− 0.25 octaves) and stronger rate decreases for tones away from CF (−0.5 to −1 octaves and above 0.5 octaves) (Fig 5a, b, c).

**Figure 5.**
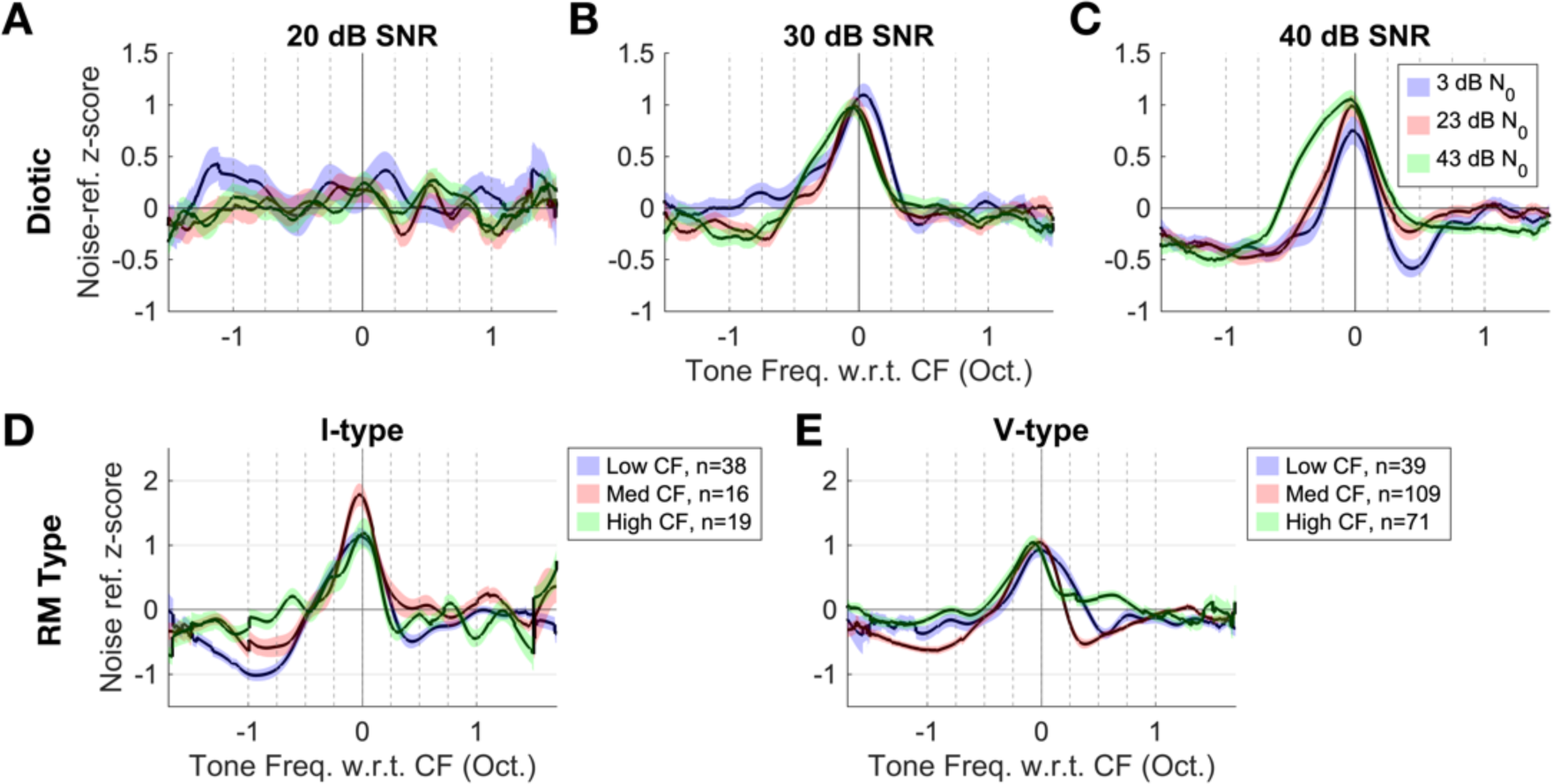
Population average responses to WB-TIN stimuli. (A-C) Diotic condition (A) 20-dB-SNR condition, (B) 30-dB SNR, (C) 40-dB SNR, blue curve is the N0 = 3-dB-SPL condition, pink curve is the N0 = 23-dB-SPL condition, and green is the N0 = 43-dB-SPL condition. All data was normalized by noise-referenced z-score. Dotted grey lines represent the tone frequencies tested in the linear mixed model. Colored bands indicate +/−1 SEM for each population average. (D-E). Population average responses to WB-TIN separated by neural characteristics. Responses averaged over contra/diotic presentation and spectrum-level presentation (3, 23, 43 dB SPL) and normalized using a noise-referenced z-score. (D) I-type and (E) V-type responses for low (blue), medium (red), and high (green) CF groups.

Changes in noise level also affected WB-TIN responses. In the 40-dB-SNR condition, increasing N_0_ broadened the excitatory peak and increased rate for tone frequencies ranging from −0.75 to +0.5 octaves relative to CF (Fig. 5c). In the 30-dB-SNR condition, when N_0_ increased, rates increased ¼ octave below CF and decreased ¼ octave above CF (Fig. 5b). No changes in rate vs. tone frequency over level were observed for the 20-dB-SNR condition, which is near threshold for TIN detection in rabbits (Zheng et al., 2002) (Fig. 5a).

The mixed-effects model also identified a significant three-way interaction between SNR, N_0_, and contralateral/diotic presentation (F(4, 20596)=7.77, p =2.95e-06) and a significant two-way interaction between tone frequency and contralateral/diotic presentation in the mixed-effects model (F_(8, 20580)=_2.15, p = 0.027); however, there were only significant differences for the tone one octave above CF. Generally, contralateral and diotic presentations of the WB-TIN stimuli resulted in similar responses (Fig. 5, Fig. S3). Overall, responses had systematic changes based on SNR and N_0_, and only the highest SNR condition revealed rate decreases below the noise-alone rate.

We wondered whether neural characteristics, such as MTF type, CF group, or RM type, changed the responses to WB-TIN. A second linear mixed-effects model was used to analyze the impact of neural characteristics. The mixed model revealed significant three-way interactions between tone frequency, RM type and CF group (F(16,16586)=3.20, p=1.45e-05, Fig. 5d, e). Low-CF I-type neurons had stronger rate decreases (−1, −0.75, 0.25 octaves w.r.t. CF) and stronger, broader rate increases (−0.25 and 0 oct) compared to low-CF V-type neurons (Fig. 5d, e, blue). At medium CFs, I-type rate increases were stronger than in V-type (0, 0.25, 0.5 oct, Fig. 5d, e, red). For high-CF neurons, V-type neurons had smaller rate decreases at 1 octave above CF (Fig. 5d, e, green). Medium-CF I-type neurons had greater rate increases (0, 0.25 oct), and weaker rate decreases (0.5) than low and high CF neurons (Fig. 5d). For V-type neurons, low CFs had broader rate increases (0.25 oct) and high CFs often had weaker or no rate decreases (−1, −0.75, 0.5 oct, Fig. 5e). A significant three-way interaction was also found between RM type, MTF type, and tone frequency (F(32,16586)=3.82, p=1.96e-12, Fig. S4). There were stronger rate increases and decreases for I-types in BE, flat, and H_BE_ neurons (Fig. S4).

Overall, RM type and CF group significantly impacted responses, and I-type neurons had stronger rate increases and decreases that varied with MTF type and CF group. Comparing CF groups, low- and mid-CF neurons had strong rate decreases off CF, but high CFs had weaker rate decreases. MTF type marginally impacted responses, suggesting that changes in the amplitude of NFs in the peripheral responses upon addition of the tone to WB noise had a relatively weak effect on these responses.

### Predicting WB-TIN responses using RMs and STRFs

We questioned to what extent the response to pure tones in quiet (RMs) could explain the WB-TIN data. If the response to WB-TIN was driven primarily by the tone, these metrics may be well correlated. To quantify similarity between RM and WB-TIN responses, we calculated the variance explained, R^2^, between a neuron’s WB-TIN response and the neuron’s RM at the sound level most closely matched to the WB-TIN stimulus. Interestingly, most V-type neurons had WB-TIN responses that were poorly explained by their RMs (43-dB-SPL N_0_, R^2^ mean = 0.24, R^2^ median = 0.17, n=121, Fig. 6a, Fig. S5a, c). I-type neurons had WB-TIN responses that were overall better explained by RMs, though there was a lot of variation in predictable variance (43-dB-SPL N_0_, R^2^ mean = 0.36, R^2^ median = 0.34, n=45, Fig. 6a, Fig. S5b, c). These results are consistent with previous reports that I-type responses to tones are robust when noise is added (Ramachandran et al., 2000). Most onset neurons RMs were not able to explain WB-TIN responses (43-dB-SPL N_0_, R^2^ mean = 0.16, R^2^ median = 0.08, n=20, Fig. S5 c). Overall, the I-type RMs were significantly more correlated to the WB-TIN response than the other two types for all sound levels (ANOVA, N_0_ = 3 dB SPL, F(2, 124)= 8.183, p=0.00046; N_0_ = 23 dB SPL, F(2, 174)=17.836, p=8.9e-8; N_0_ = 43 dB SPL, F(2, 181)=9.202, p=0.00016). However, variance explained by the RM alone was generally poor.

**Figure 6.**
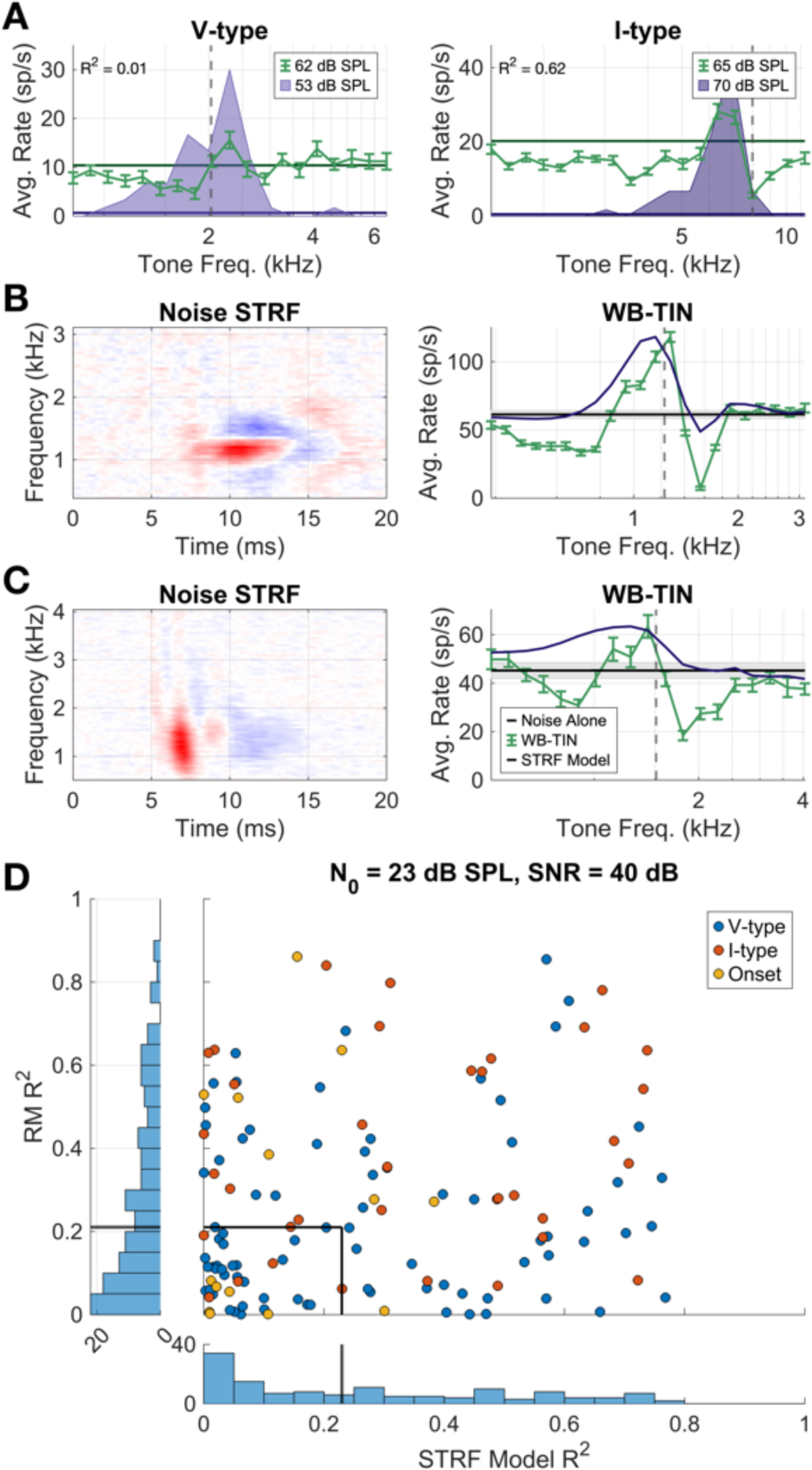
Example STRF predictions of WB-TIN and summary of predictions based on RM and STRF. (A) Response map (purple) with WB-TIN response (green) overlayed for two example neurons, one V-type (left) and one I-type (right). (B) Example neuron STRF (left) and STRF predictions of WB-TIN (right, purple) compared to actual WB-TIN response (right, green), R^2^ = 0.66. (C) Different example neuron for same analysis as (B), R^2^ = 0.20. (D) Scatter plot of RM R^2^ and STRF R2 values. Grey lines represent the median R^2^ values.

Another approach to characterizing neurons is to use an STRF, which is a linear mapping of firing rate to stimulus features, used to describe spectral and temporal tuning properties of IC neurons (Escabí and Schreiner, 2002; Andoni et al., 2007). Many IC STRFs reveal excitatory and inhibitory regions at varied frequencies and delays, which could be related to the WB-TIN responses. Here we compare WB-TIN responses with a prediction using a noise STRF because both stimuli use gaussian noise. We tested whether STRFs could predict the responses to WB-TIN for the 23-dB-SPL N_0_ condition. STRF model predictions of WB-TIN responses were compared to the WB-TIN data using the *R*^2^ goodness-of-fit metric. Results varied; most WB-TIN responses were not well predicted by the STRF, but 25/145 neurons had a *R*^2^ > 0.5. One such neuron had an STRF with excitation near CF and delayed on-CF and off-CF inhibition (Fig. 6b). The STRF model predicted the neural WB-TIN response well, with the largest errors in the size and frequency range of the inhibition (*R*^2^ = 0.66, Fig. 6b). A second example neuron that was not well predicted had an STRF with a broad excitation and broad, delayed inhibition (Fig. 6c). The STRF prediction did not accurately reflect the inhibition seen in WB-TIN responses (R^2^=0.20 Fig. 6c). There was no correlation between the RM *R*^2^ and the STRF-model *R*^2^ for the 23-dB-SPL N_0_ conditions and no correction for specific RM types (Fig. 6d). Overall, STRF and RMs were not sufficient to explain the WB-TIN responses, but STRFs provided the best predictions for neurons that had STRFs with inhibitory sideband regions.

### Difference of gaussians (DoG) fit to WB-TIN data

Noting that the STRF predictions with inhibitory sidebands tended to have more accurate WB-TIN predictions, we next asked how broad inhibition factored into these WB-TIN responses. A DoG function was fit to each average-rate WB-TIN profile to quantify patterns in response profiles across neurons for the diotic, N_0_= 23 dB SPL, 40-dB-SNR condition (Fig 7). An example fit including the excitatory and inhibitory components is shown in Fig. 7a. The DoG fit well (variance explained, R^2^ > 0.5) to the majority of neurons that had high predictable variances (Fig. 7b, blue). The ratio of inhibition to excitation amplitude was generally below one (mean = 0.92, median = 0.81), meaning that inhibition was weaker than excitation in the model fit for most neurons (Fig. 7c). The DoG equation had parameters for both the center frequency of the excitatory gaussian (𝑓_𝑒𝑥𝑐_) and the inhibitory gaussian (𝑓_𝑖𝑛ℎ_) to allow fits to asymmetrical off-CF inhibition. The inhibitory and excitatory center frequencies were similar to the CF of the neuron but were often lower than CF (Fig. 7d). The difference between inhibitory and excitatory center frequencies had a mean of 0.197 octaves with respect to CF, indicating on average the inhibitory CF skewed to lower frequencies compared to the excitatory CF and created stronger lower inhibitory sidebands. The bandwidths of excitation (𝜎_𝑒𝑥𝑐_) and inhibition (𝜎_𝑖𝑛ℎ_) were plotted as a function of CF and fit using a log-log transformed linear model (Fig. 7e). Bandwidths for both excitation (Fig. 7e, top) and inhibition (Fig. 7e, middle) increased as a function of CF (p<0.001 for all excitatory conditions, p=0.002 for inhibition, 23-dB-SPL N_0_, Fig. 7b). For many units, inhibitory bandwidths were wider than excitatory bandwidths (n=129/192, Fig. 7e, bottom). The same trends were seen in N_0_= 3 dB SPL and N_0_= 43 dB SPL, 40-db-SNR condition (Fig. S6). Scatter plot shows each model fit as a function of log bandwidth vs. log strength ratio, with example profiles for each quadrant (Fig 7f). Model fits in quadrant one have negative rate profiles due to strong inhibition strength and broad inhibition (n=22/192). Fits in quadrant two can have two profiles: strong, narrow inhibition can fall outside of the excitatory gaussian and thus represent an excitation with a single sideband of inhibition (n=17/192) or can represent a positive rate profile with a dip near CF (n=22/192). Quadrant three, representing weak inhibition and narrow inhibition, can have a profile with a single inhibitory sideband (n=24/192). Lastly, quadrant four, strong excitation with broad inhibition, represents the “classic” difference of gaussians fit with excitation near CF and broad, weaker inhibition off CF. Quadrant four contains 55.7% of the fits and represents the majority of the data (n=107/192). Points near (0,0) represent profiles that were generally flat, where inhibition and excitation were nearly equal in strength and bandwidth. Overall, the majority of neural responses to WB-TIN were successfully fit using a difference of gaussians and the majority of these neurons had broad, weak inhibition, but there was heterogeneity in the data and other rate profiles were seen as well.

**Figure 7.**
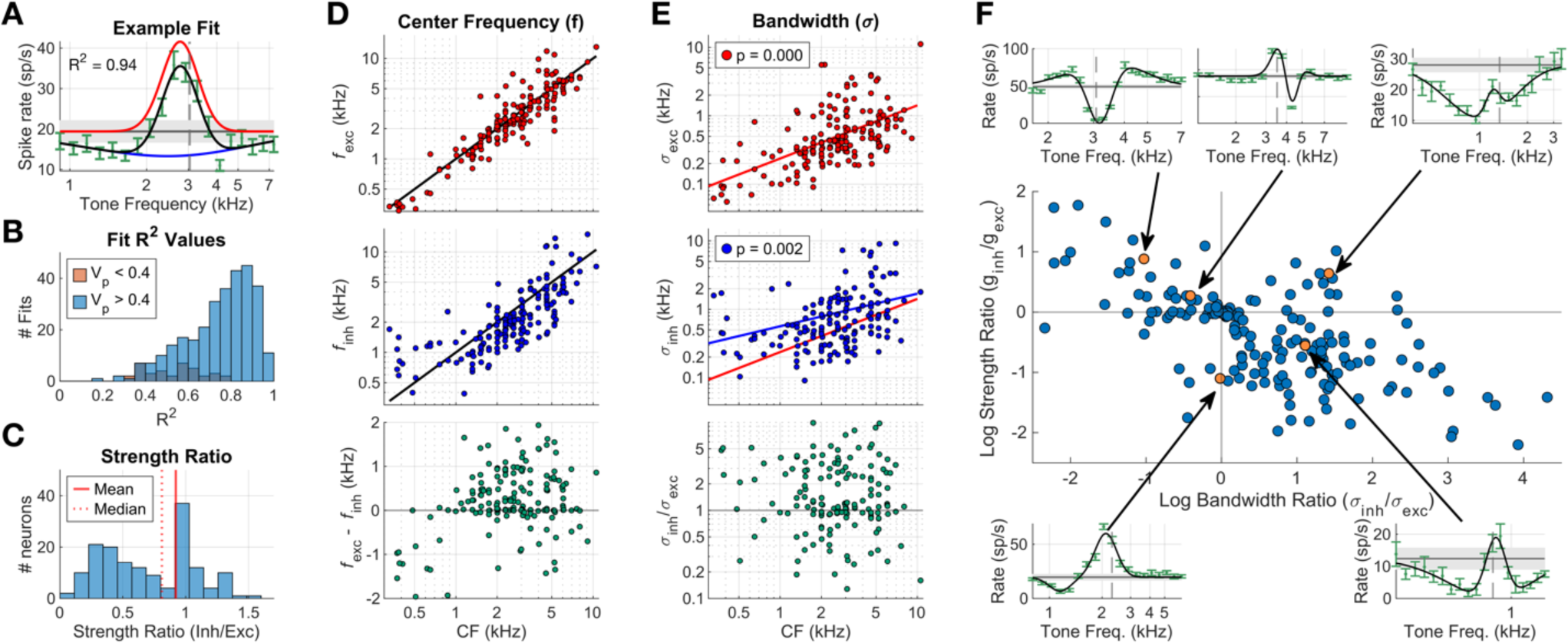
Difference of gaussians analysis. (A) DoG fit to an example neuron 40-dB SNR WB-TIN response. Red curve is the excitatory gaussian, blue curve is the inhibitory gaussian. Green bars represent the data and +/−1 SEM. Black curve is the DoG. Horizontal black line indicates the noise-alone rate, and grey shaded area is +/− 1 SEM. (B) Histogram of R^2^ values fit to all neurons in the diotic, 23 dB SPL, 40-dB SNR condition. Orange indicates neurons with a predictable variance less than 0.4. (C) Strength ratio (inh/exc) for all fit neurons. Red solid line indicates the mean, red dotted line indicates the median. (D) Center frequencies for excitatory (red, top) and inhibitory (blue, middle) gaussians as a function of CF. Difference between excitatory and inhibitory center frequencies in green, bottom row. (E) Bandwidth, 𝜎, for excitatory (red, top) and inhibitory (blue, middle) gaussians. σ_𝑒𝑥𝑐_fit lines in red (top, middle), σ_𝑖𝑛ℎ_fit line in blue (middle). Bottom row, green, indicates the ratio of inhibitory to excitatory bandwidths. (F) Scatter of the strength ratio vs bandwidth ratio with example neuron fit curves shown for each quadrant. Orange points indicate examples.

### Adding off-CF pathways to IC model improved model accuracy

Lastly, we added off-CF inhibitory pathways to an existing IC SFIE model with an on-CF excitation and a delayed inhibition (Nelson and Carney, 2004; Carney and McDonough, 2019). The goal was to implement the broad inhibition seen in the DoG fits to predict WB-TIN responses in a phenomenological model of the IC, while also retaining amplitude-modulation sensitivity, a key feature of IC neurons. Six model configurations with off-CF inhibitory inputs from different sources (brainstem, BS IC cell, BE IC cell) targeting on-CF IC neurons (BS or BE) were tested to search for model configurations that could replicate physiological results (Fig. 2).

Two exemplar neurons were used to fit the model configurations, one BE cell and one BS cell (Fig. 8a, Fig 9a). Although the WB-TIN results were not affected by amplitude-modulation sensitivity, on-CF NB-TIN results depend on MTF type. Thus, we investigated whether these model configurations could retain accurate MTFs and NB-TIN responses while also fitting the WB-TIN rate profiles.

**Figure 8.**
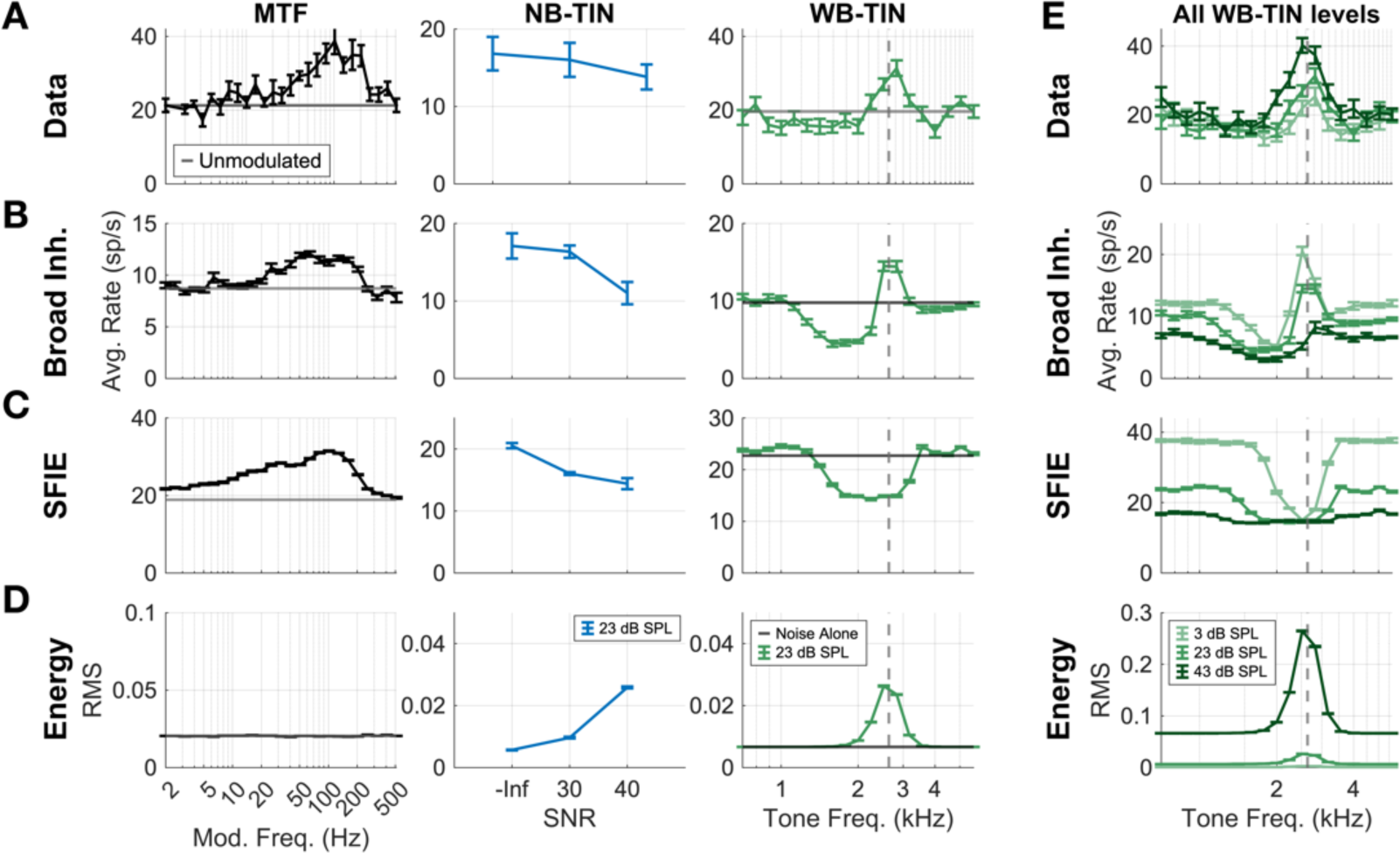
Comparison of model responses to data for three models: broad inhibition, SFIE, and energy. (A) Example BE neuron MTF (left), NB-TIN response (middle), and WB-TIN response for the N_0_=23 dB SPL, SNR = 40 dB condition (overall level = 50, 62 dB SPL respectively). These three datasets were used for fitting the broad inhibition model. (B) Broad inhibition model responses to MTF, NB-TIN, WB-TIN. S_low_ = 1, S_high_= 0.28, D_off_ = 1 ms, off-CF range = 0.625 octaves, BMFs = [87 164 111] for [low-CF on-CF high-CF]. (C) SFIE model responses. (D) Energy model responses. (E) Data, broad inhibition, SFIE, and energy model responses for three levels of WB-TIN (3, 23, 43 dB SPL, 40 dB SNR). Error bars represent +/−1 SEM.

**Figure 9.**
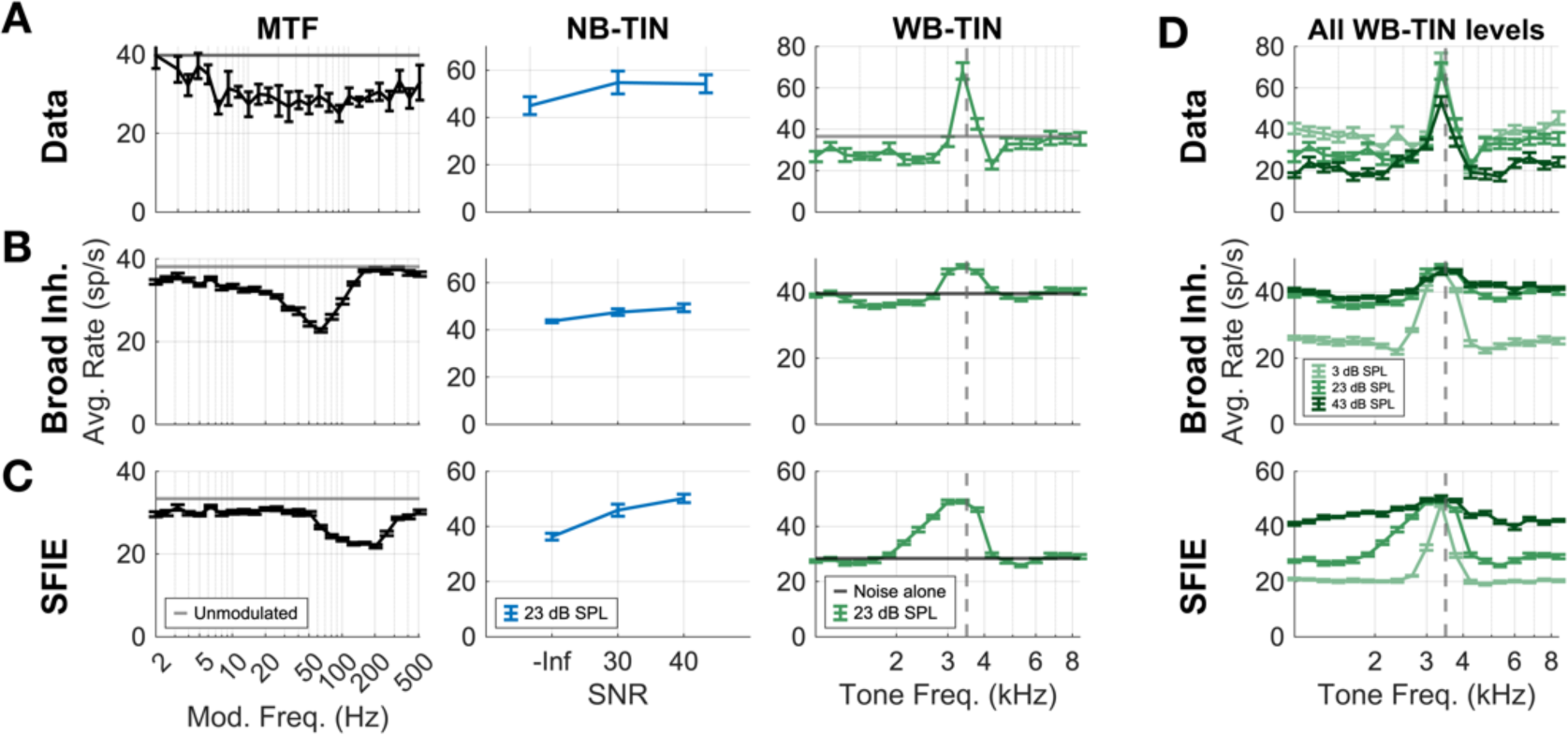
Comparison of model responses to data for three models: broad inhibition, SFIE, and energy. (A) Example BS neuron MTF (left), NB-TIN response (middle), and WB-TIN response for the N_0_=23 dB SPL, SNR = 40 dB condition (overall level = 50, 62 dB SPL respectively). These three datasets were used for fitting the broad inhibition model. (B) Broad inhibition model responses to MTF, NB-TIN, WB-TIN. S_low_ = 0.19, S_high_= 0, D_off_ = 1 ms, off-CF range = 0.5 octaves, BMFs = [300 38 10] for [low-CF on-CF high-CF]. octaves (C) SFIE model responses. (D) Data, broad inhibition, SFIE, and energy model responses for three levels of WB-TIN (N_0_=3, 23, 43 dB SPL, 40 dB SNR). Error bars represent +/−1 SEM.

The model without added inhibition was tested as a baseline. The SFIE BE model had decreased rate in response to on-CF NB-TIN and predicted a dip in rate at CF for the WB-TIN stimulus, which poorly predicted the neural data (r_MTF_ =0.78; r_NB_ = 0.86; r_WBTIN_=0.03, Fig. 8c). The energy model was also tested as a baseline, and as expected, predicted peaks in rate when the tone frequency matched CF but did not predict the rate decreases below and above CF (r_MTF_ = −0.34; r_NB_ = −1.00; r_WBTIN_=0.56, Fig. 8d). All six model configurations were fit to this BE cell; the BS-inhibited-by-BS model configuration (Fig. 2c) was able to replicate the MTF, on-CF NB-TIN, and WB-TIN results and MTF type with the best accuracy (r_MTF_ = 0.81; r_NB_ = 0.99; r_WBTIN_=0.85, Fig. 8b).

All models tested had limitations when predicting responses over a range of sound levels. For the energy model (Fig. 8e, bottom), the increase in rate when the tone was off-CF was too large. The response of the SFIE model (Fig. 8e) to noise-alone or to a noise with a tone far from CF decreased as level increased, in stark contrast to the neural data, due to the model sensitivity to neural fluctuations in the AN model responses. As sound level increased, even for a noise, the fluctuations in the AN responses decrease due to IHC saturation, and the SFIE BE cell responded to a lack of fluctuations with a decrease in rate. The response to a tone at CF was also not correct over a wide range of sound levels for the SFIE and broad inhibition models. The tone captured the AN response and decreased fluctuations, thus decreasing the BE SFIE rate, but reached a ‘floor’ when the AN response was fully captured. The broad inhibition model decreased in rate as level increased near CF, due to the broadening of inhibitory pathways as sound level increased. The broadening inhibition decreased excitation at CF.

The exemplar BS response was fit using a model BS neuron that was inhibited by two off-CF BS neurons (Fig. 2c, Fig. 9). Again, the SFIE BS cell without lateral inhibition was a baseline. The SFIE BS model increased in rate with increasing SNR for the on-CF NB-TIN stimulus, and there was a peak in rate at CF for the WB-TIN response. This model accurately predicted the on-CF rate increases, but not the off-CF rate decreases seen in data (r_MTF_ = 0.39; r_NB_ = 0.94; r_WBTIN_=0.39, Fig. 9c). The SFIE BS WB-TIN response featured a small amount of inhibition on the high-frequency side due to suppression of the AN responses. However, that inhibition was not strong enough to accurately model the physiological IC responses. With the additional off-CF BS inhibitory inputs to the on-CF BS cell, inhibition was stronger than with the on-CF pathways alone, and this model had the greatest accuracy (r_MTF_ = 0.53; r_NB_ = 0.93; r_WBTIN_=0.52, Fig 9b). Similar to the BE SFIE and broad-inhibition models, these BS models also did not accurately predict changes in level (Fig 9d). These inaccuracies were created again by the SFIE model’s response to decreases in fluctuations to noise as level increased, causing an increase in rate for noise-alone and off-CF tone responses in the BS models not seen in physiology.

The other configurations tested that did not accurately predict WB-TIN were the three models with an on-CF BE target cell and the BS-inhibited-by-off-CF-BE model (not shown). The model configurations that feature an on-CF BE target cell always decreased in rate near CF due to the underlying on-CF BE cell, which responded to WB-TIN with a dip near CF. Adding off-CF inhibition to the BE-model configuration caused a larger decrease in rate. As the strength of inhibition was increased, the rates further decreased to zero. The BS-inhibited-by-off-CF-BE model configuration also did not replicate inhibition off CF in the IC responses in any combination of model parameters tested. The off-CF BE cell responded with a decrease in rate when the tone was near the CF, which lead to the on-CF BS cell being inhibited by a weak off-CF response. This model configuration resulted in a lack of inhibition off CF in the WB-TIN response. Lastly, the BS-inhibited-by-off-CF-CN model did not predict the off-CF inhibition observed in the IC responses.

An interesting aspect of the broad-inhibition models is the way BE and BS cells were created from only a combination of BS cells (Fig 10), largely due to BMF and inhibitory strength parameters. To illustrate these MTFs, the on- and off-CF BS cell responses were plotted individually (Fig 10, top row). The on-CF BS cell for the BE model fit has a shallow BS MTF, and the off-CF MTFs have more salient BS MTFs (Fig 10b, top). When combined, the on-CF response is strongly inhibited at low and high modulation frequencies, and less inhibited at the WMF of the off-CF response (approx. 100 Hz), resulting in a broad-inhibition model with a BE MTF profile (Fig 10b, bottom). The broad-inhibition BS cell had a stronger BS on-CF MTF that was weakly inhibited by the off-CF BS cells, which were flatter due to changes in WMF parameters (Fig 10c).

**Figure 10.**
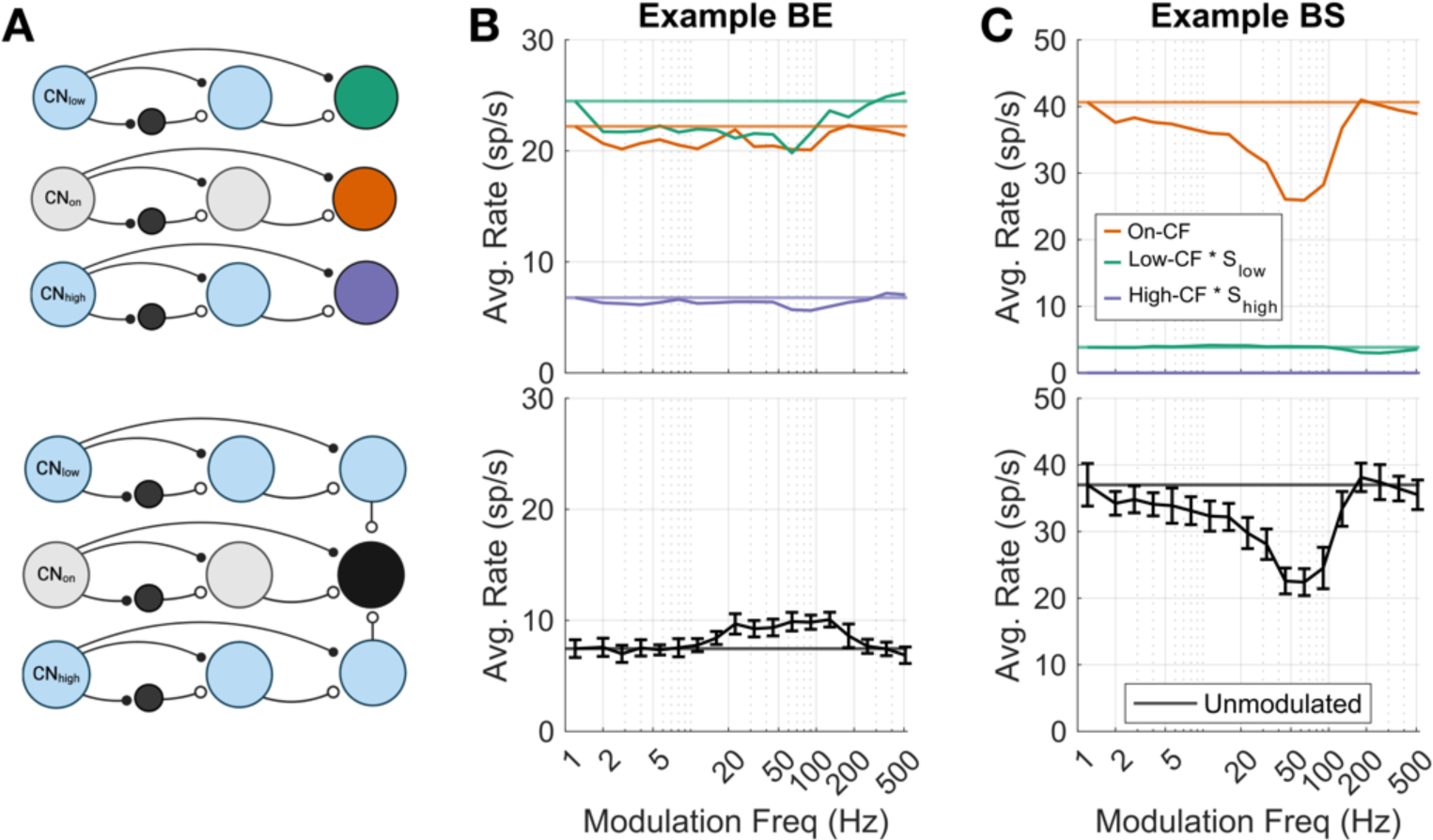
Model MTF responses for intermediate components of the broad inhibition model. (A) Model schematics, top row has on-CF (orange), low-CF (green), and high-CF (purple) cells highlighted. Bottom row shows full broad inhibition model cell (black). (B) Example BE broad inhibition model response, top row has on-CF, low-CF, and high-CF MTFs plotted separately with the low- and high-CF responses scaled by inhibition strength, same parameters as Fig 8. Bottom row is the aggregate broad inhibition response. (C) Same plots for an example BS broad inhibition cell, for Fig 9 model cell.

We further investigated how changing the parameters in the model would influence model responses by varying the strengths, delays, on- and off-CF parameters. Varying the strength (S_off_), or weight of the inhibitory inputs, resulted in changing MTF types and changing overall rate responses to WB-TIN (Fig. 11a). At S_off_ = 0.1 the MTF is BS, at 0.3 the MTF is hybrid, and at 0.5 the MTF is BE (Fig. 11a). As S_off_ increases, the overall rate response to WB-TIN decreased and inhibitory regions became more salient (Fig. 11a). Next, we changed the CF of the off-CF inputs and found that, as expected, the further away in frequency the off-CF inputs were, the broader the inhibitory regions of the response were (Fig. 11b). The MTF was not largely impacted by changing the CF-range parameter (Fig. 11b). We also tested off-CF delays, 𝐷_𝑜𝑓𝑓_, of 0, 1, and 2 ms and found this parameter had a slight impact on the MTF of the neuron but had no effect on the WB-TIN response (not shown). Lastly, the CF of the on-CF model neuron was varied to compare WB-TIN and MTF responses across a range of CFs (Fig. 11c). The 1000-Hz model neuron with S_off_ = 0.4, 𝐷_𝑜𝑓𝑓_ = 0 ms, and off-CF range = 1 octave created a mostly flat MTF with a WB-TIN response with excitation and flanking inhibition (Fig. 11c). The 3-kHz model neuron, which was considered a ‘mid’ CF range, with the same model parameters, resulted in a hybrid MTF and WB-TIN responses that were consistent with physiological responses. The high-CF 5-kHz model neuron had a BE MTF, and the WB-TIN responses did not have as clear excitation and inhibition (Fig. 11c). Overall, adding an off-CF input from local BS model cells to the on-CF BS cell (Fig. 4c) created responses that retained MTF sensitivity by replicating BE, BS, hybrid, and flat MTF types and accurately predicted WB-TIN responses seen in physiological recordings.

**Figure 11.**
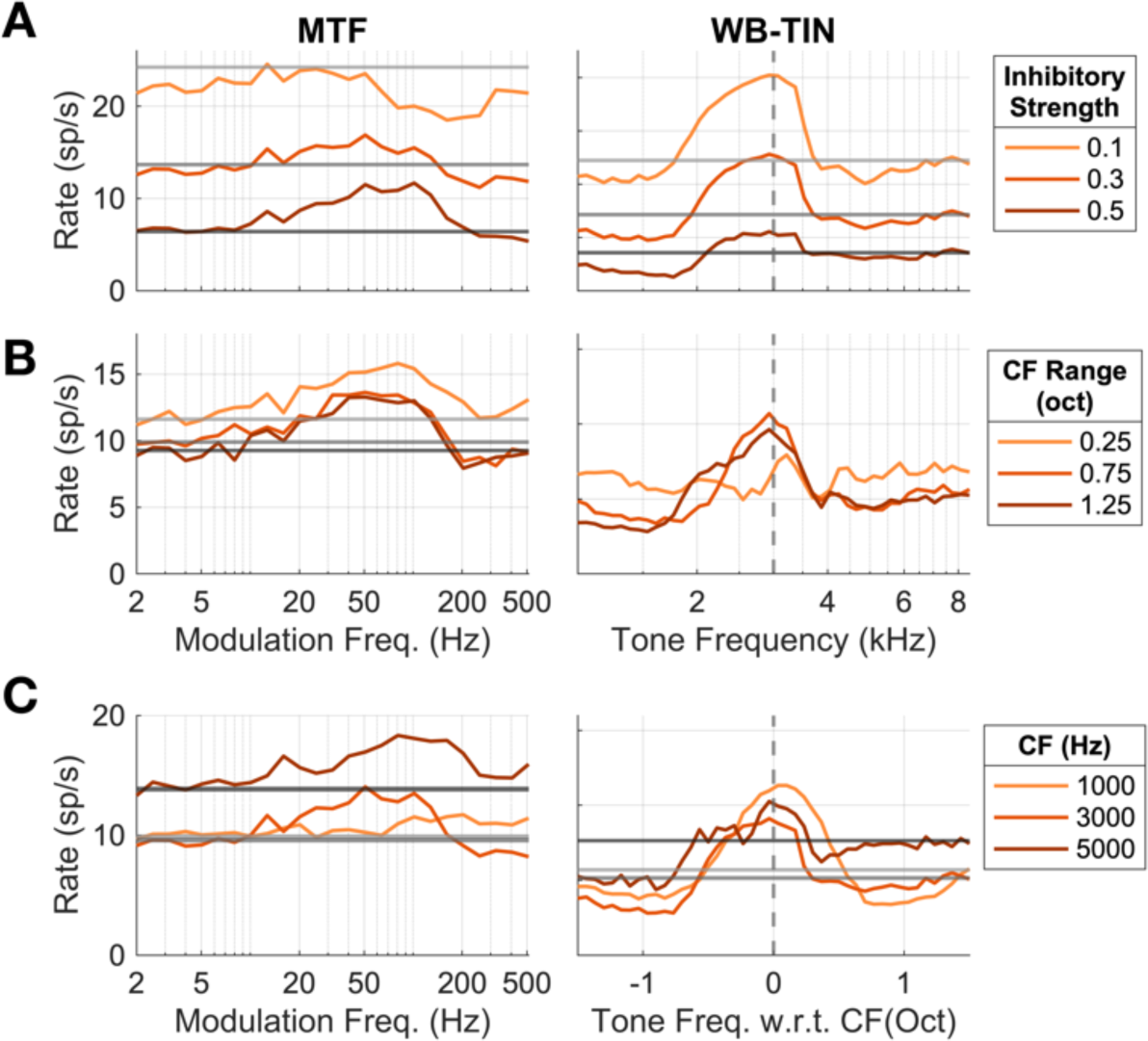
Varying lateral-inhibition model (see Fig. 4H) parameters changes model responses. (A) Lateral SFIE model response for a 3 kHz model neuron with S_off_ parameter set to 0.1 (light orange), 0.3 (orange), or 0.5 (brown). D_off_ = 0 ms, off-CF range = 1 octave. Model responses to MTF and WB-TIN with N_0_ = 23 dB SPL. Noise-alone responses for WB-TIN shown in light grey (0.1), grey (0.3) and dark grey (0.5). (B) Responses of a 3-kHz model neuron with off-CF range parameter set to 0.25 (light orange), 0.75 (orange), or 1.25 (brown) octaves away from CF. S_off_ = 0.4, D_off_ = 0 ms. Noise-alone response not shown for clarity. (C) Responses of a 1-kHz (light orange), 3-kHz (orange), and 5-kHz (brown) model neurons with S_off_ = 0.4, D_off_ = 0 ms, off-CF range = 1 octave. All models have BMF = 100 Hz.

## 4. Discussion

Our study revealed that average rates in response to WB-TIN increased for tones near CF and decreased for tones away from CF, compared to responses to noise-alone stimuli. This pattern differed from responses to narrowband noise, suggesting that mechanisms in addition to NF sensitivity impact TIN encoding. This pattern also often differed from what would be expected based on neurons’ tone RMs and noise STRFs. The responses to WB-TIN featured off-CF rate decreases that occurred for a range of sound levels and at suprathreshold SNRs, supporting the hypothesis that broad inhibition shaped responses. We developed a model that incorporated on-CF excitation and inhibition with two off-CF inhibitory inputs from local BS model cells that accurately predicted WB-TIN responses while maintaining MTF sensitivity. Overall, these results highlight broad inhibition as a mechanism influencing complex-sound encoding in the IC.

Our approach of shifting the tone frequency with respect to CF can also be used to infer the population response of IC neurons. In this inferred population, the neurons tuned near the tone frequency are excited and the neurons tuned away from the tone frequency are inhibited with respect to their noise-alone response. This combination of excitation and inhibition would create a rate profile across a population of neurons with enhanced representation of a spectral peak in noise.

### Comparisons to TIN IC studies

Our findings are consistent with studies that observed rates that increased with SNR for an on-CF tone in wideband noise (Rees and Palmer, 1988; Jiang et al., 1997; Rocchi and Ramachandran, 2018). For on-CF NB-TIN, BE IC neural responses were largely consistent with Fan et al. (2021), showing decreased rates with increasing SNR. However, BS cell responses here were more variable and did not consistently show increased rates with increasing SNR (Fig. 4). A possible explanation could be methodological: we used diotic MTFs instead of contralateral MTFs and did not rely on identification of CF during the experiment because we sampled many tone frequencies. We observed that IC responses to NB-TIN often changed from a rate increase to a rate decrease at different tone frequencies near CF, highlighting the impact of the choice of CF value to test (not shown).

IC responses to off-CF tones have not been widely studied, though one group found that responses to off-CF tones helped to explain behavioral TIN-detection thresholds in budgerigar (Wang et al., 2021). That finding, combined with our results that off-CF rates contain information in the form of inhibition, supports the idea that off-CF information could be important in TIN detection.

We originally hypothesized that ipsilateral inhibition in the binaural response may strengthen off-CF inhibitory bands in responses, but overall did not see changes in inhibitory sidebands due to presentation ear in our population. However, differences in diotic and contralateral responses on an individual neuron level could be studied in tandem with interaural response properties in the future.

### Broad Inhibition

Broad inhibition may arise from various sources, including GABAergic projections from the contralateral and ipsilateral dorsal nucleus of the lateral lemniscus (DNLL) (Adams and Mugnaini, 1984), glycinergic input from the ipsilateral lateral superior olive and ventral nucleus of the lateral lemniscus (Saint-Marie et al., 1989; Saint-Marie and Baker, 1990; Winer et al., 1995), or local or commissural connections within the IC itself (Saldaña and Merchań, 1992). There is also evidence of lateral inhibition or suppression in earlier auditory areas such as ventral and dorsal cochlear nucleus (Martin and Dickson, 1983; Rhode and Greenberg, 1994).

Lateral inhibition is also postulated to enhance the rate representation of spectral information at high sound levels (Shamma, 1985). Our physiological results showed that inhibition was more pronounced at high SNRs (40 dB SNR w.r.t. No) than low SNRs (30 dB SNR), consistent with reports that STRFs at higher sound levels have increased areas of inhibition (Lesica and Grothe, 2008). These results suggests that strength of inhibition may be level or SNR dependent, which could contribute to sound processing in the IC at high levels. Future physiological work understanding the origin and function of broad inhibition in the subcortical auditory system is necessary to understand the impact of broad inhibition.

### Modeling

Our broad-inhibition computational model is phenomenological and thus replicates IC responses, not physiological structure. However, we do think the model is physiologically possible. Firstly, in our model fitting process we found that local inhibition of BS cells was the only configuration that could replicate IC responses, consistent with physiological results that found blocking local IC inhibitory receptors reduced off-CF inhibition in STRFs (Andoni et al., 2007).

Secondly, the model relies on two subpopulations of BE cells, because the on-CF BS cell was created from an on-CF BE cell, which is a limitation currently (Fig 2c). However, we hypothesize that the initial BE stage may represent neurons in other regions, such as the DNLL. The DNLL receives input from the ventral CN and provides strong inhibitory input to IC cells. Additionally, some neurons in the DNLL have band-pass rate MTFs (Yang and Pollak, 1997). Therefore, the first BE cell stage can represent the DNLL.

Lastly, we found that as the WB-TIN level was increased, the overall excitation decreased in model BE cells, contrary to physiological responses. We explored adding efferent feedback (Farhadi et al., 2023), but this model, which was designed based on narrowband stimuli, did not improve the predictions of our wideband responses. The incorrect responses over level were due to saturated on- and off-CF HSR AN fibers that provided the ascending inputs to the model IC neuron. A new IC model is needed to overcome this issue, possibly by including wideband efferent feedback pathways.

### Alternative Explanations

Suppression and efferent feedback are alternative mechanisms that could appear as off-CF inhibition in WB-TIN responses. Suppression occurs when a tone outside of the tuning curve of an AN fiber decreases the rate of the fiber in response to a tone at CF (Abbas and Sachs, 1976; Delgutte, 1990). In response to WB-TIN, AN suppression could result in a rate profile in the IC that appears to have inhibitory sidebands. However, the AN model used replicates suppression and the BS SFIE model response did not have decreased rates for tones below CF and had smaller decreases above CF as compared to physiological data (Fig. 9c), suggesting that suppression alone cannot account for the changes in rate in WB-TIN responses.

Subcortical efferent feedback also has the potential to alter responses in the IC. Responses in the IC to AM stimuli increase or decrease over a timescale consistent with the medial olivocochlear (MOC) efferent system, dependent on MTF type, indicative that efferent feedback may change responses in the IC (Farhadi et al., 2023). Many neurons adapted over time in response to WB-TIN, and some neurons increased in rate over duration for specific tone frequencies (not shown), possibly due to efferent effects. Additionally, some MOC neurons project to outer hair cells across many frequency channels (Brown, 2014). This wide-bandwidth efferent feedback, which could impact IC responses to off-CF tones, is a topic for future study.

### Visual retinal ganglion cell & DoG models

The excitatory region and inhibitory sidebands seen in the WB-TIN data are reminiscent of retinal ganglion cell (RGC) responses in the visual system. RGCs have center-surround organization, with excitation (or inhibition, for off-center RGCs) when light is within a receptive field, and inhibition (or excitation) when light is presented in the surrounding area (Rodieck, 1965). This mechanism arises from lateral inhibition and enhances edge and contrast detection in a visual scene (Boycott and Wässle, 1999). Our IC results show that a tone is presented at a frequency near CF excited neurons, but tones away from CF neuron inhibited or suppressed responses. We hypothesize that the excitation-inhibition pattern in IC cell responses to TIN serves to increase contrast, making the tone response a more salient feature in the rate code.

DoG models, previously used for RGCs (Rodieck and Stone, 1965; Enrorh-Cugell and Robson, 1984) were used to fit WB-TIN responses. Additionally, DoG spectral receptive field models fit to IC rate responses to harmonic complex tones predicted responses more accurately than a gaussian due to the added inhibition (Su and Delgutte, 2020). Our results had similar parameter fit trends to the Su and Delgutte study, with one exception: we observed a strong correlation between inhibitory bandwidth and CF, which they did not report.

### Future Experiments / Summary

The comparison in this study between NF sensitivity and broad inhibition raises the following question: at what bandwidth does inhibition become more prevalent than NF sensitivity? Future studies could include a band-widening experiment to test the bandwidth-dependence of NF-sensitivity or inhibition mechanisms. The shifting tone frequency in a bed of noise is also similar to the two-tone paradigm, which also reveals inhibitory regions (Egorova et al., 2001). It would be interesting to know whether a response map created using the two-tone paradigm more accurately reflects WB-TIN responses. Lastly, an STRF created using ripples may result in better predictions than noise STRFs, as these ripple STRFs perform better for higher auditory areas (Andoni et al., 2007; Versnel et al., 2009).

In conclusion, this study revealed distinct response patterns of rate increases to on-CF tones and rate decreases to off-CF tones added to broadband noise in the IC. These results suggest that broad inhibition may have a critical role in complex-sound encoding by shaping rate profiles of sounds in a noisy background. A computational model of the IC including both broad inhibition and AM sensitivity could explain both narrowband and wideband TIN results, highlighting the complexity in encoding different TIN stimuli. These findings could impact our understanding of how natural sounds with broad bandwidths and amplitude modulation, such as speech and music, are represented in the rate profile of IC neurons.

## Acknowledgements

This study was funded by NIH-R01-DC010813 and NIH-F31-DC020630-03. Thanks to Paul W. Mitchell and Swapna Agarwalla for assisting in data collection, and to Douglas Schwarz for assistance with hardware and software. Thanks to Kris Abrams for helping with the rabbit surgery and care.

