## Supplemental for "Mechanisms of Tone-in-Noise Encoding in the Inferior Colliculus"

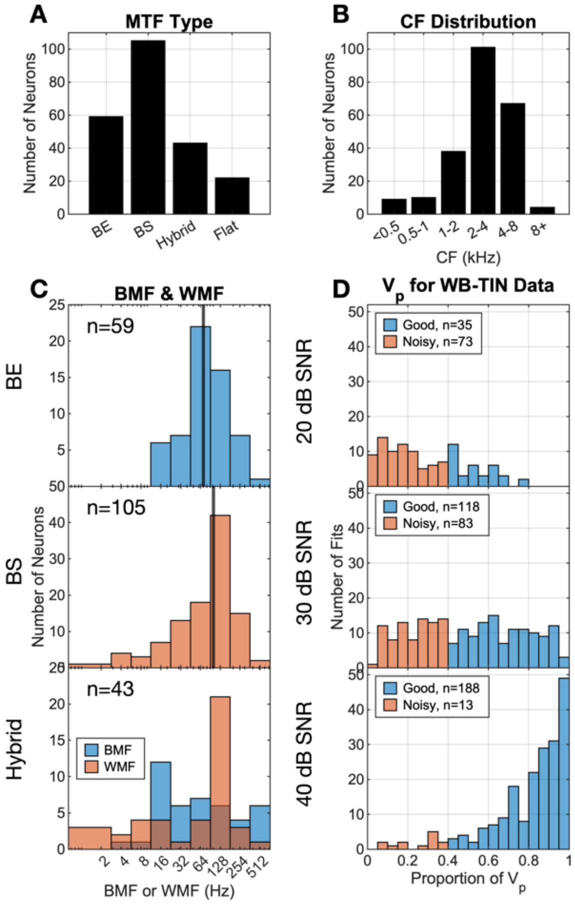


**Supplemental 1.** CF and MTF distributions of neurons, n = 229. (A) Distributions for each MTF type. (B) CF distribution of dataset, in octaves. (C) Distribution of BMFs for BE neurons, median = 70 Hz. Distribution of WMFs for BS neurons, median = 102 Hz. Distributions of BMFs and WMFs for hybrid neurons, BMF median = 64 Hz, WMF median = 125 Hz. (D) Predictable variance distributions for 20-, 30-, and 40-dB SNR for binaural $N_0$= 23dB SPL dataset.

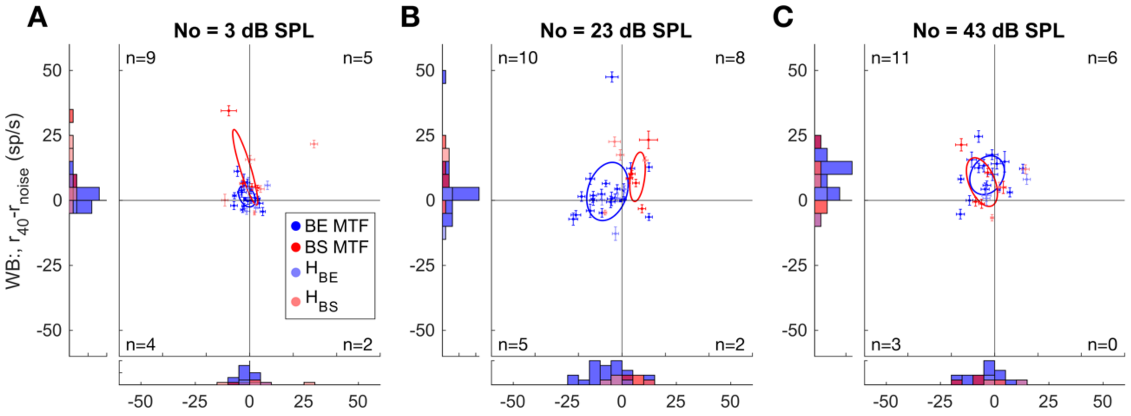


**Supplemental 2.** Differences between average rates in response to contralateral TIN (40 dB SNR) and noise-alone for on-CF tones in wideband vs. narrowband noise. Responses at (A) N0 = 3 dB SPL, WB-TIN overall level range of 34-48 dB SPL, NB-TIN level range of 28-41 dB SPL, (B) 23 dB SPL, WB-TIN overall level range of 54-69 dB SPL, NB-TIN level range of 48-63 dB SPL, and (C) 43 dB SPL, WB-TIN overall level range of 74-83 dB SPL, NB-TIN level range of 68-83 dB SPL. BE MTFs are blue, BS MTFs are red, HBE are light blue, HBS are pink. Error bars represent +/- 1 SEM. Ellipses represent the standard deviation for BE (blue) and BS (red) neurons. Number of neurons in each quadrant is reported in the corner of each quadrant.

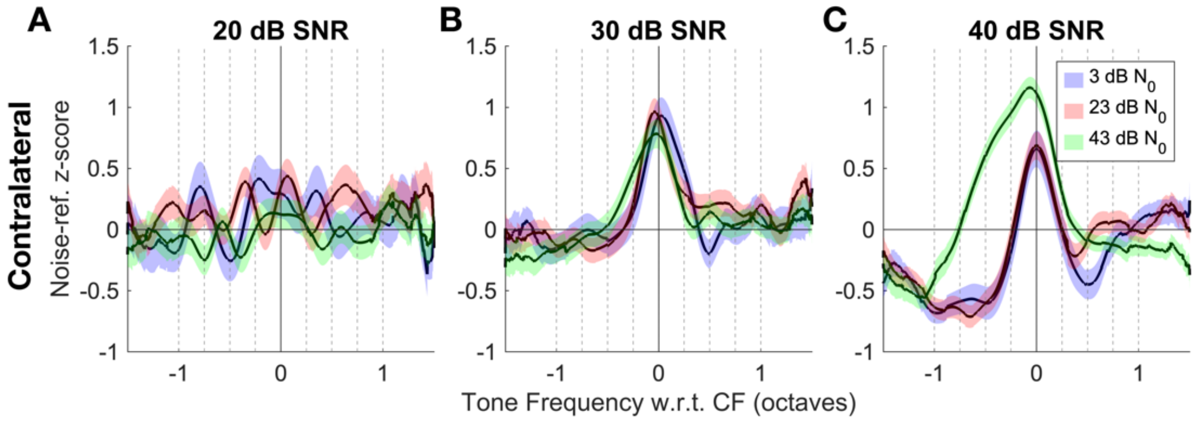

**Supplemental 3.** Population average responses to contralateral WB-TIN stimuli. (A) 20-dB-SNR condition, (B) 30-dB-SNR, (C) 40-dB-SNR, blue curve is the $N_0$ = 3-dB-SPL condition, pink curve is the $N_0$ = 23-dB-SPL condition, and green is the $N_0$ = 43-dB-SPL condition. All data was normalized by noise-referenced z-score. Dotted grey lines represent the tone frequencies tested in the linear mixed model. Colored bands indicate +/-1 SEM for each population average.

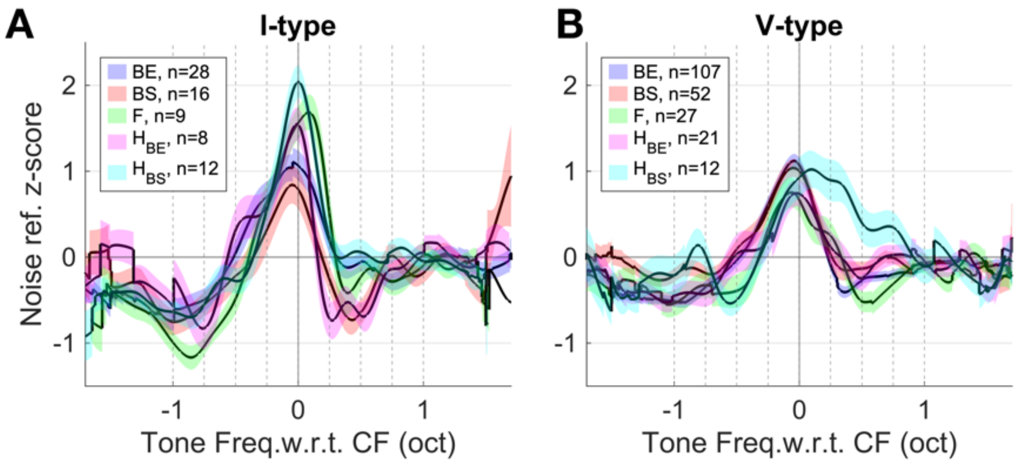


**Supplemental 4.** Population average responses to WB-TIN separated by neural characteristics. Responses averaged over contra/diotic presentation and spectrum-level presentation (3, 23, 43 dB SPL, 40-dB-SNR condition) and normalized using a noise-referenced z-score. (A) I-type and (B) V-type responses for BE (blue), BS (red), flat (green), HBS (pink), and HBE (cyan). Dotted grey lines represent the tone frequencies tested in the linear mixed model. Colored bands indicate +/-1 SEM for each population average.

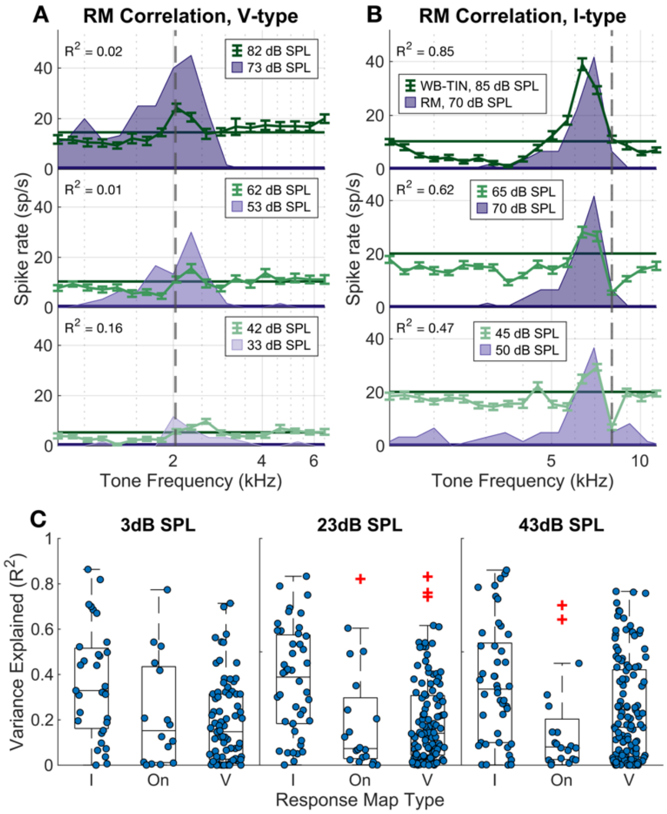


**Supplemental 5.** Variance explained in the WB-TIN responses by neuron responses to pure tones for diotic stimuli. (A) Example V-type unit, CF = 2036 Hz, pure tone responses at 33, 53, and 73 dB SPL in purple. Response to WB-TIN at three levels, $N_0$ = 3, 23, 43 dB SPL in green. (B) Example I-type neuron, presentation, CF = 8000 Hz. (C-E) Variance explained for all units, condition. Neurons were split into three groups: V-type, I-type, and onset.

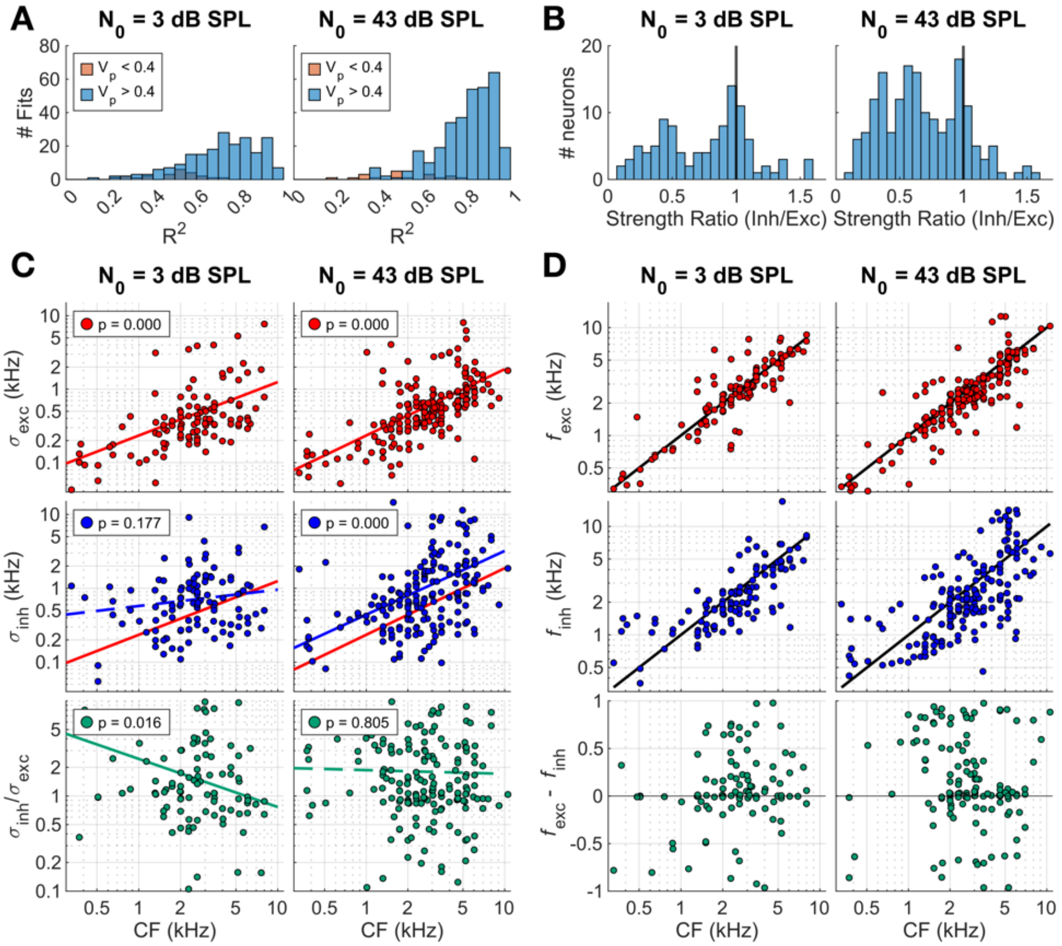


**Supplemental 6.** Difference of gaussians analysis for 3- and 43-dB SPL. (A) Histogram of $R^2$ values fit to all neurons in the diotic, 40-dB SNR, 3 dB SPL (left) and 23 dB SPL (right) condition. Orange indicates neurons with a predictable variance less than 0.4. (B) Strength ratio (inh/exc) for all fit neurons. (C) Bandwidth, $\sigma$, for excitatory (red, top) and inhibitory (blue, middle) gaussians. $\sigma_{exc}$ fit lines in red (top, middle), $\sigma_{inh}$ fit line in blue (middle). Bottom row, green, indicates the ratio of inhibitory to excitatory bandwidths. (D) Center frequencies for excitatory (red, top) and inhibitory (blue, middle) gaussians as a function of CF. Difference between excitatory and inhibitory center frequencies in green, bottom row.
